# Tofacitinib uptake by patient-derived intestinal organoids predicts individual clinical responsiveness

**DOI:** 10.1101/2024.03.02.583137

**Authors:** Kyung Ku Jang, Defne Ercelen, Jing Yu Carolina Cen Feng, Sakteesh Gurunathan, Chaoting Zhou, Aryeh Korman, Luke Newell, David Hudesman, Drew R. Jones, P’ng Loke, Jordan E. Axelrad, Ken Cadwell

## Abstract

**Background & Aims:** Despite increasing therapeutic options in the treatment of ulcerative colitis (UC), achieving disease remission remains a major clinical challenge. Nonresponse to therapy is common and clinicians have little guidance in selecting the optimal therapy for an individual patient. This study examined whether patient-derived materials could predict individual clinical responsiveness to the Janus kinase (JAK) inhibitor, tofacitinib, prior to treatment initiation.

**Method:** In 48 patients with UC initiating tofacitinib, we longitudinally collected clinical covariates, stool, and colonic biopsies to analyze the microbiota, transcriptome, and exome variations associated with clinical responsiveness at week 24. We established patient-derived organoids (n = 23) to determine how their viability upon stimulation with proinflammatory cytokines in the presence of tofacitinib related to drug responsiveness in patients. We performed additional biochemical analyses of organoids and primary tissues to identify the mechanism underlying differential tofacitinib sensitivity.

**Results:** The composition of the gut microbiota, rectal transcriptome, inflammatory biomarkers, and exome variations were indistinguishable among UC patients prior to tofacitinib treatment. However, a subset of patient-derived organoids displayed reduced sensitivity to tofacitinib as determined by the ability of the drug to inhibit STAT1 phosphorylation and loss of viability upon cytokine stimulation. Remarkably, sensitivity of organoids to tofacitinib predicted individual clinical patient responsiveness. Reduced responsiveness to tofacitinib was associated with decreased levels of the cationic transporter MATE1, which mediates tofacitinib uptake.

**Conclusions:** Patient-derived intestinal organoids predict and identify mechanisms of individual tofacitinib responsiveness in UC. Specifically, MATE1 expression predicted clinical response to tofacitinib.

## INTRODUCTION

Ulcerative colitis (UC), a subtype of inflammatory bowel disease (IBD), is a chronic inflammatory disorder of the gastrointestinal tract that affects millions of individuals worldwide^1, 2^. Although etiology of UC is unknown, the leading hypothesis is that disease results from combinations of microbial and environmental factors in genetically susceptible individuals^1, 2^. There is no cure for UC, and medical therapies are aimed at controlling symptoms, promoting remission, and preventing relapse^3, 4^. Despite increasing availability of immunomodulatory treatments, disease relapse is common.^5^ Many patients are resistant to a given biologic or small molecule [primary nonresponders (NRPs)] and a significant proportion become refractory after an initial improvement (secondary NRPs)^3, 6, 7^. Thus, predicting drug responsiveness prior to therapy initiation may provide critical guidance to clinicians in optimizing therapy selection for individual patients and improving outcomes.

Tofacitinib is an oral small molecule indicated for patients with moderate to severe UC refractory to anti-tumor necrosis factor (TNF) therapy. Tofacitinib inhibits activation of Janus kinase (JAK) proteins to prevent phosphorylation of signal transducer and activator of transcription (STAT) that is required for transcription of inflammatory genes downstream of cytokine receptors^8^. Tofacitinib preferentially inhibits JAK1 and JAK3 signaling, and to a lesser extent JAK2^9^. Despite the increasing utilization of JAK inhibitors, primary and secondary nonresponse occurs in more than 30% and 25% of patients, respectively^10–13^. Although several reports have analyzed clinical data, gut microbiota, and mucosal transcriptomes to identify distinguishing factors associated with responsiveness to inhibitors of TNFα and α_4_β_7_ integrins^14–17^, there are little data addressing JAK inhibitors.

The recent emergence of organoid technology has provided a reliable *in vitro* model for IBD research^18–20^. Patient-derived intestinal organoids (PDIOs) represent a 3D culture system that recreates the architecture and cell composition of an individual’s *in vivo* intestinal epithelium by culturing stem cell-containing crypts seeded into a supporting matrix with media containing growth factors^18^. As they retain genetic and epigenetic alterations of their originating tissue, PDIOs have been applied to study phenotypic variance. PDIOs reproduce epithelial abnormalities associated with an *ATG16L1* variant and somatic mutations in inflamed epithelia of UC patients^21–23^. Furthermore, PDIOs maintain abnormalities of DNA methylation in IBD patient-derived intestinal epithelia^24, 25^. Consequently, PDIOs display heterogeneity in response to cytokines, microbes, anti-cancer drugs, and therapies for cystic fibrosis^21, 23, 26–29^, suggesting that PDIOs can be applied to predict individual drug responsiveness, achieving precision medicine for UC patients.

In this study, we collected patient-derived materials and clinical metadata to determine the factors distinguishing RPs and NRPs to tofacitinib. Less than half of patients responded to tofacitinib, consistent with real-world experience data^12^. RPs and NRPs were indistinguishable by analyses of the gut microbiota, rectal transcriptome, inflammatory biomarkers, clinical metadata, and exome variants prior to tofacitinib treatment. We developed an *ex vivo* platform to recreate an individual’s IBD-related microenvironment from intestinal biopsies by culturing PDIOs with inflammatory cytokines and assessed the ability of tofacitinib to inhibit cytokine-induced cell death. RP-derived organoids displayed improved viability when treated with tofacitinib, which was associated with higher expression of the cationic transporter multidrug and toxin extrusion protein 1 (MATE1) that mediates tofacitinib uptake. JAK inhibitors that are independent of MATE1-mediated uptake rescued viability of tofacitinib-insensitive organoids. Our findings reveal a mechanism in which patients with UC are primarily unresponsive to tofacitinib using PDIOs.

## METHOD

### Sample and data collection from cohort receiving tofacitinib

We enrolled and followed patients initiating tofacitinib for moderate to severe UC at the Inflammatory Bowel Disease Center at New York University (NYU) Langone Health between January 1, 2019 and June 30, 2022. Demographic and UC-related clinical data were collected at baseline and during follow up at week (W)0, 2, 12, and 24, including the partial and total Mayo score^30^. Stool samples were collected from patients at W0, 2, 12, and 24. Blood samples were collected at W0. Endoscopic pinch biopsies from the rectum and ascending colon were obtained at W0 and 24. In addition, endoscopic pinch biopsies were obtained from ascending colon from 4 healthy adult control subjects undergoing screening colonoscopy. The inflammation status of tissue was confirmed by expert IBD histopathological examination.

Subject information is shown in Supplementary Tables 1-3. Over the study period, patients were categorized by clinical response to tofacitinib as RPs, partial responders (PRPs), or NRPs. An RP was defined as a patient with a decrease from the W0 baseline total Mayo score of at least 3 points and at least 30%, with an accompanying decrease in the rectal bleeding subscore of at least 1 point or an absolute rectal bleeding subscore of 0 or 1 with persistence on tofacitinib at W24. A PRP was defined as a patient with a decrease from W0 baseline total Mayo score < 3 points and < 30% with persistence on tofacitinib at W24. An NRP was defined as a patient with no change from W0 baseline total Mayo score at W24 and/or discontinuation of tofacitinib during the study period due to lack of clinical effect.

This protocol was approved by the NYU Grossman School of Medicine Institutional Review Board (Mucosal Immune Profiling in Patients with Inflammatory Bowel Disease, S12-01137 and Identify Predictors that Distinguish Between Tofacitinib Responders and Non-responders Based on Genotype and Cellular and Molecular Profiles from Pinch Biopsies, Blood, and Stool Samples, S18-00630), and all participants provided informed consent. The clinical data were collected using EPIC EHR and REDCap 9.3.6 software.

### DNA extraction,16S ribosomal RNA (rRNA) sequencing analysis, and fecal calprotectin measurement

DNA from human stool samples were extracted with DNeasy PowerSoil Pro kit (Qiagen) according to the manufacturer’s instruction. Bacterial 16S rRNA genes were amplified at the V4 region using primer pairs and sequenced by Illumina MiSeq system at the NYU Genome Technology Core. Sequencing reads were processed using the DADA2 pipeline in the QIIME2 software package. Taxonomic assignment was performed against the Greengenes 13_8 99% OTUs full-length sequences database^31^. Alpha diversity analysis was done using observed OTUs^32^. Beta diversity was calculated using Bray-Curtis, Jaccard, unweighted UniFrac, and weighted UniFrac distance and visualized with EMPeror^33^. Quantification of fecal calprotectin was performed using Calprotectin ELISA Assay Kit (Eagle Biosciences) according to the manufacturer’s instructions.

### RNA-sequencing (RNA-seq)

Total RNA was extracted from 48 rectal pinch biopsies that were collected at W0 or W24 (Supplementary Figure 3A) using RNeasy Mini Kit with DNase treatment (Qiagen) according to the manufacturer’s instructions. The libraries generated using TruSeq Standard Total RNA Library Prep with RiboZero Gold rRNA removal kit (Illumina) were sequenced by NovaSeq 6000 (Illumina) at NYU Genome Technology Core. RNA-seq results were processed using the R package DESeq2 to obtain variance stabilized count reads, fold changes relative to specific condition, and statistical *P* value. Analysis of the rectal transcriptome focused on differentially expressed genes (DEGs), defined as the genes with an absolute fold change relative to specific condition >2 and an adjusted *P* value < 0.05. Enriched pathways, upstream regulators, and cytokine for the DEGs were analyzed by ingenuity pathway analysis (Qiagen). The analyses were visualized using R packages ggplot2 and ComplexHeatmap^34^.

### Exome sequencing

Genomic DNA from blood samples was purified using QIAamp DNA Blood Kits (Qiagen) according to the manufacturer’s instructions and sequenced by Psomagen using SureSelect XT Human All Exon V8 kit (Agilent) on NovaSeq 6000. Whole exome fastq files were processed using Genome Analysis Toolkit Best Practices^35^ and annotated using Variant Effect Predictor^36^. Variants that showed a significant difference in allele frequency between IBD patients and healthy controls from 1000 Genomes Project^37^ were identified and used for association analyses with Enrichr^38, 39^. Furthermore, the Mutect2 and GenomicsDBImport tools from GATK were used to create a panel from RPs. This panel and population controls from the Genome Aggregation Database^40^ were compared to NRPs to identify variants that may explain differences between tofacitinib response groups.

### Culture of human colonic organoids

Colonic organoids were cultured as described previously^26, 27^. Endoscopic mucosal pinch biopsies were collected in ice-cold complete Roswell Park Memorial Institute (RPMI) [RPMI 1640 medium supplemented with 10% fetal bovine serum, 100 IU penicillin and 100 μg/ml streptomycin (Corning), 2 mM L-glutamine (Corning), and 50 μM 2-mercaptoethanol (ThermoFisher)]. The pinch biopsies were incubated in Gentle Cell Dissociation Reagent (Stemcell Technologies) on ice for 30 min followed by vigorous pipetting to isolate crypts. The crypts were embedded in 30 μl of Matrigel and cultured with Human IntestiCult Organoid Growth Medium (Stemcell Technologies), herein referred to as maintenance media. The culture medium was changed every 2-3 days. For passing organoids, 10 μM Y-27632 (Sigma-Aldrich) was added for the first 2 days.

### Viability assay of human colonic organoids

Mature colonic organoids grown in the maintenance media were mechanically disrupted by vigorous pipetting and suspended thoroughly in 30 μl Matrigel at approximately 5-7 debris/μl. For differentiation, the maintenance media was replaced with advanced DMEM/F-12 (ThermoFisher) in the presence of 100 IU penicillin and 100 μg/ml streptomycin, 125 μg/ml gentamicin (ThermoFisher), 2 mM L-glutamine, 20 ng/ml epidermal growth factor (EGF) (PeproTech), 100 ng/ml Noggin (R&D Systems), 1 μg/ml R-spondin 1 (R&D Systems), 100 ng/ml rWnt-3a (R&D systems), 1 mM N-acetylcysteine (Sigma-Aldrich), 10 nM gastrin (Sigma-Aldrich), 500 nM A-83-01 (Tocris), 200 ng/ml insulin-like growth factor 1 (IGF-1) (Biolegend), 100 ng/ml fibrobalst growth factor 2 (FGF-2) (Peprotech), and 1x B27 (ThermoFisher), herein referred as IGF-1 and FGF-2 (IF) medium^41^. Each 10 μl drop was cultured in a 96 -well culture plate in triplicates with IF medium supplement with10 μM of Y-27632 for the first 2 days. Organoids were washed 3 times using advanced DMEM/F12 to remove Y-27632 and proceeded to the viability assay.

For treatment with JAK inhibitors, tofacitinib (pan JAK inhibitor), upadacitinib (JAK1 inhibitor), fedratinib (JAK2 inhibitor), and ruxolitinib (JAK1/2 inhibitor) were introduced to organoids (Supplementary figure 10D). Briefly, organoids were cultured in the IF medium supplemented with TNFα (20 ng/ml, R&D Systems) and interferon (IFN)β (1,000 IU/ml, R&D Systems) or TNFα (20 ng/ml) and IFNγ (20 ng/ml, R&D Systems) in the absence or presence of tofacitinib (1 or 10 μM, Pfizer), upadacitinib (0.1 or 1 μM, Selleckchem), fedratinib (0.1 or 1 μM, Selleckchem), or ruxolitinib (0.1 or 1 μM, Selleckchem) for another 4 days.

For treatment with MATE1-selective inhibitors, ritonavir (1, 5, 10, or 20 μM, Sigma-Aldrich), famotidine (1, 10, or 20 μM, Sigma-Aldrich), or indinavir (1, 10, or 20 μM, Sigma-Aldrich) were treated with the organoids stimulated with cytokines±JAK inhibitors for another 4 days. An organic cation transporter (OCT)2-selective inhibitor pantoprazole (1, 10, or 20 μM, Sigma-Aldrich) was used as a negative control (Supplementary figure 10A). The culture medium was changed every 3 days. Percent viable organoids were determined by quantification of the number of intact organoids on days 0 and 4 as descried previously^21^. Opaque organoids with condensed structures or those that have lost adherence were excluded from viability assessment.

### Transcript analysis

Total RNA was extracted from the colonic organoids using RNeasy Mini Kit with DNase treatment (Qiagen) according to the manufacturer’s instruction. Synthesis of complementary DNA (cDNA) was conducted with High-Capacity cDNA Reverse Transcription Kit (ThermoFisher) according to the manufacturer’s protocol. Real time (RT)-PCR was performed using SybrGreen (Roche) on a LightCycler 480 system (Roche) using the following primers: *JAK1*, Fwd 5’-CCAGAACTGCCCAAGGACAT-3’ and Rev 5’-CGCATCCTGGTGAGAAGGTT-3’; *STAT1*, Fwd 5’-CTTACCCAGAATGCCCTGAT-3’ and Rev 5’-CGAACTTGCTGCAGACTCTC-3’; *TNFα*, Fwd 5’-CCAGGCAGTCAGATCATCTTCTC-3’ and Rev 5’-AGCTTGAGGGTTTGCTACAACAT-3’; *ISG15*, Fwd 5’-GAGAGGCAGCGAACTCATCT-3’ and Rev 5’-CTTCAGCTCTGACACCGA CA-3’; *INFβ1*, Fwd 5’-GCCGCATTGACCATCTAT-3’ and Rev 5’-GTCTCATTCCAGCCAGTG-3’; *IFNλ2/3*, Fwd 5’-GCCACATAGCCCAGTTCAAG-3’ and Rev 5’-TGGGAGAGGATATGGTGCAG-3’; *OCTN1*, Fwd 5’-CTACATCGTCATGGGTAGTC-3’ and Rev 5’-CCAGATCTGAACCATTTCAC-3’; *OCTN2*, Fwd 5’-GACCATATCAGTGGGCTATTT-3’ and Rev 5’-CTGCATGAAGAGAAGGACAC-3’; *MATE1*, Fwd 5’-ATGCTGTTTCCCACCTCTTTG-3’ and Rev 5’-TCCAACCTTCTGATTTCCACTC-3’; *MATE2*, Fwd 5’-GCAGGGCCAGTTTTCATTTA-3’ and Rev 5’-TGGGAGATGATGTTGGCATA-3’; *GAPDH*, Fwd 5’-GATGGGATTTCCATTGATGACA-3’ and Rev 5’-CCACCCATGGCAAATTCC-3’. The relative expression of the respective genes to *GAPDH* expression was calculated using the ΔΔ*C*_T_ method^42^ and was expressed as fold change normalized to unstimulated organoids.

### Western blotting

Colonic organoids were processed for immunoblotting as previously described^27^. Briefly, the organoids were washed with phosphate-buffered saline (PBS), incubated with Cell Recovery Solution (Corning) at 4°C for 30 min to dissociate Matrigel, and centrifuged at 400 x *g* for 5 min. The pellets were suspended in lysis buffer [20 mM Tris-HCl (pH 7.4, NYU Reagent Preparation Core), 150 mM NaCl (NYU Reagent Preparation Core), 1% Triton X-100 (Sigma-Aldrich), 10% glycerol (Sigma-Aldrich), and 2× Halt Protease and Phosphatase Inhibitor Cocktail (ThermoFisher)] and pelleted twice at 10,000 x *g* for 10 min at 4°C to collect the lysates. The organoid lysates were resolved on Bolt 4-12% Bis-Tris Plus Gels (Invitrogen), transferred onto polyvinylidene difluoride membranes, and blocked using Intercept (TBS) blocking buffer (LI-COR). Membranes were probed with primary antibody overnight at 4°C. The following primary antibodies were used for western blotting studies: anti-STAT1 (Cell Signaling, 9176S), Phospho-STAT1 (Tyr701) (Cell Signaling, 9167S), JAK1 (Cell Signaling, 3332S), MATE1 (Cell Signaling, 14550S), organic cation transporter novel (OCTN)1/2 (Santa Cruz, sc-515731), and β-actin (Sigma-Aldrich, A5441). After incubation with the primary antibody, the membrane was washed and probed with the secondary antibody for 1 hr at room temperature. As for secondary antibodies, IRDye 680RD Goat anti-Rabbit (925-68071) and IRDye 800CW Goat anti-Mouse (925-32210) were purchased from LI-COR. After additional washing, protein was detected with Image Studio for Odyssey CLx (LI-COR). Band intensities were measured by Fiji/ImageJ.

### Liquid chromatography/mass spectrometer (LC/MS)

To measure extracellular concertation of tofacitinib, mature organoids (around 1,600) grown in maintenance media were mechanically disrupted and suspended thoroughly in 200 μl Matrigel. Each 25 μl drop was cultured in a 24-well culture plate with IF medium supplement with10 μM of Y-27632 for 2 days. Organoids were then washed 3 times using advanced DMEM/F12 to remove Y-27632 and then cultured with IF medium for another 5 days. On day 5, the organoids were treated with 10 μM of tofacitinib for 4 hr and culture supernatants were collected and stored at - 80°C.

The supernatants mixed with 100% methanol (ThermoFisher) and 625 nM amino acid standard cocktail (Research Products International) were homogenized with disruption beads (Research Products International) using bead blaster homogenizer (Benchmark Scientific) to extract metabolites. Samples were centrifuged at 21,000 *x g* for 3 min at 4°C. The metabolite supernatants were dried down by SpeedVac Vacuum Concentrator (ThermoFisher) and reconstituted in LC/MS grade ethanol (Sigma-Aldrich), sonicated, and centrifuged to collect the metabolite lysates. A matrix-controlled 7-point standard curve of tofacitinib (0.01, 0.03, 0.1, 0.3, 1, 3, and 10 μM) was prepared, extracted, and analyzed as technical duplicates alongside the supernatant samples.

The samples were subjected to an LC/MS analysis to quantify extracellular concentration of tofacitinib. The LC/MS parameters were adapted from a previously described method^43^. Injection volume was set to 2 μl for all analyses The samples were separated using LC column (Dionex Ultimate 3000 system (ThermoFisher) coupled with Water BEH-Phenyl (2.1 × 150 mm, 1.7 μm) (Waters)). The column oven temperature was set to 25°C for the gradient elution. A flow rate of 200 μl/min was used with the following buffers; A) 0.1% formic acid in water, and B) 0.1% formic acid in acetonitrile. The gradient profile was as follows; 0-35% B (0-10 min), 35-75% B (10-15 min), 75-99% B (15-15.25 min), 99-99% B (15.25-16.5 min), 99-0% B (16.5-16.75 min), and 0-0% B (16.75-20 min).

MS analyses were carried out by coupling the LC system to a Thermo Q Exactive HF mass spectrometer (ThermoFisher) operating in heated electrospray ionization mode. Method duration was 20 min with a polarity switching data-dependent Top 5 method for both positive and negative modes. Spray voltage for both positive and negative modes was 3.5 kV and capillary temperature was set to 320°C with a sheath gas rate of 35, aux gas of 10, and max spray current of 100 μA. The full MS scan for both polarities utilized 120,000 resolution with an automatic gain control (AGC) target of 3 × 10^6^ and a maximum injection time (IT) of 100 ms, and the scan range was from 95-1000 m/z. Tandem MS spectra for both positive and negative mode used a resolution of 15,000, AGC target of 1 × 10^5^, maximum IT of 50 ms, isolation window of 0.4 m/z, isolation offset of 0.1 m/z, fixed first mass of 50 m/z, and 3-way multiplexed normalized collision energies of 10, 35, and 80. The minimum AGC target was 1 × 10^4^ with an intensity threshold of 2 × 10^5^. All data were acquired in profile mode.

### Immunofluorescence

MATE1 staining was performed as described previously^27, 44^. Briefly, mucosal pinch biopsies of the ascending colon were fixed in 10% formalin and embedded in paraffin blocks. Sections were cut to 5 μm thickness at the NYU Center for Biospecimen Research and Development and mounted on frosted glass slides. For deparaffinization, slides were baked at 70°C for 1.5 hr, followed by rehydration in descending concentration of ethanol (100%, 95%, 80%, 70%, and ddH_2_O twice; each step for 30 sec). Heat-induced epitope retrieval was performed in a pressure cooker (Biocare Medical) using Dako Target Retrieval Solution, pH 9 (Dako Agilent) at 97°C for 10 min and cooled down to 65°C. After further cooling to room temperature for 20 min, slides were washed for 10 min 3 times in Tris-buffered saline (TBS) containing 0.1% Tween 20 (Sigma-Aldrich, TBS-T). Sections were blocked in 5% normal donkey serum (Sigma-Aldrich) in TBS-T at room temperature for 1 hr, followed by incubation with rabbit anti-MATE1 antibody (1:500) in the blocking solution at 4°C overnight. Sections were washed 3 times with TBS-T and stained with the Alexa Flour 555 conjugated with donkey anti-rabbit IgG (1:500, ThermoFisher, A-31572) in PBS with 3% bovine serum albumin (BSA) (ThermoFisher), 0.4% saponin, and 0.02% sodium azide at room temperature for 1 hr. Following this, sections were washed 3 times with TBS-T and mounted with ProLong Glass Antifade Mountant with NucBlue Stain (ThermoFisher). Images were acquired using an EVOS FL Auto Cell Imaging System (ThermoFisher) and then processed and quantified using Fiji/ImageJ.

### Quantification and statistical analysis

Bars in the figure panels represent mean values ± standard error of mean. Statistical methods in this study are described in the figure legend using GraphPad Prism 9. N represents individual patient or individual organoid line described in the figure legend. *P* < 0.05 is considered statistically significant for all assays, and individual *P* values are indicated in the figure panels.

## RESULTS

### Patient population

We enrolled 48 patients initiating tofacitinib for UC. At baseline (W0), the mean age was 34.7 years (range 17-68), 54% were male (n = 26), and mean duration of UC was 9.5 years (range 2-40; Supplementary Table 1). 68.5% (n = 33), 29.2% (n = 14), and 2.1% (n = 1) had pancolitis, left-sided colitis, and proctitis, respectively. All patients had prior exposure to a biologic therapy. At baseline, the mean fecal calprotectin was 3115.5 µg/g (range 78-21169) with 97.9% (n = 47) demonstrating a Mayo endoscopic subscore of 2 or 3. At W24, 37.5% (n = 18) comprised RPs, 56.3% (n = 27) were either PRPs or NRPs, and 6.3% (n = 3) were lost to follow up (Supplementary Table 2). Stratifying by response at W24, RPs, PRPs, and NRPs had similar disease activities at W0.

### Characterization of gut microbiota in tofacitinib-treated UC patients

Previous reports suggested microbial and genetic factors are associated with anti-TNFα responsiveness^14, 15, 45^. To identify microbial factors associated with tofacitinib responsiveness, we performed 16S rRNA sequencing of stool samples collected at W0, 2, 12, and 24 (Supplementary Figure 1A). However, the microbiota compositions among RPs, PRPs, and NRPs were similar according to alpha and beta diversities at W0, 2, 12, or 24 (Supplementary Figures 1B-E and 2A-D). Also, alpha and beta diversities of RPs, PRPs, and NRPs were unchanged during tofacitinib treatment (Supplementary Figures 1F-H and 2E-G).

### Rectal transcriptome is comparable among UC patients at W0 but diverges according to tofacitinib responsiveness at W24

We analyzed the transcriptome of rectal biopsies collected at W0 and W24 (Supplementary Figure 3A). The number of transcripts displaying > 2-fold changes (adjusted *P* < 0.05) in 9 pairwise comparisons are summarized in Figure 1A. The condition that displayed the most differences was that comparing NRPs to RPs at W24. Pairwise comparisons among RPs, PRPs, and NRPs on W0 displayed no differences, indicating that the transcriptomes according to tofacitinib responsiveness are indistinguishable at W0. When comparing W0 with W24, 68 and 20 genes were differentially expressed in RPs and PRPs, respectively (Figure 1A), indicating that tofacitinib influences the rectal transcriptomes in RPs and PRPs but not in NRPs. Principal component analysis (PCA) showed that transcriptomes did not segregate based on drug responsiveness at W0, but that RPs separated from NRPs at W24 (Supplementary Figure 3B and C).

**Figure 1.**
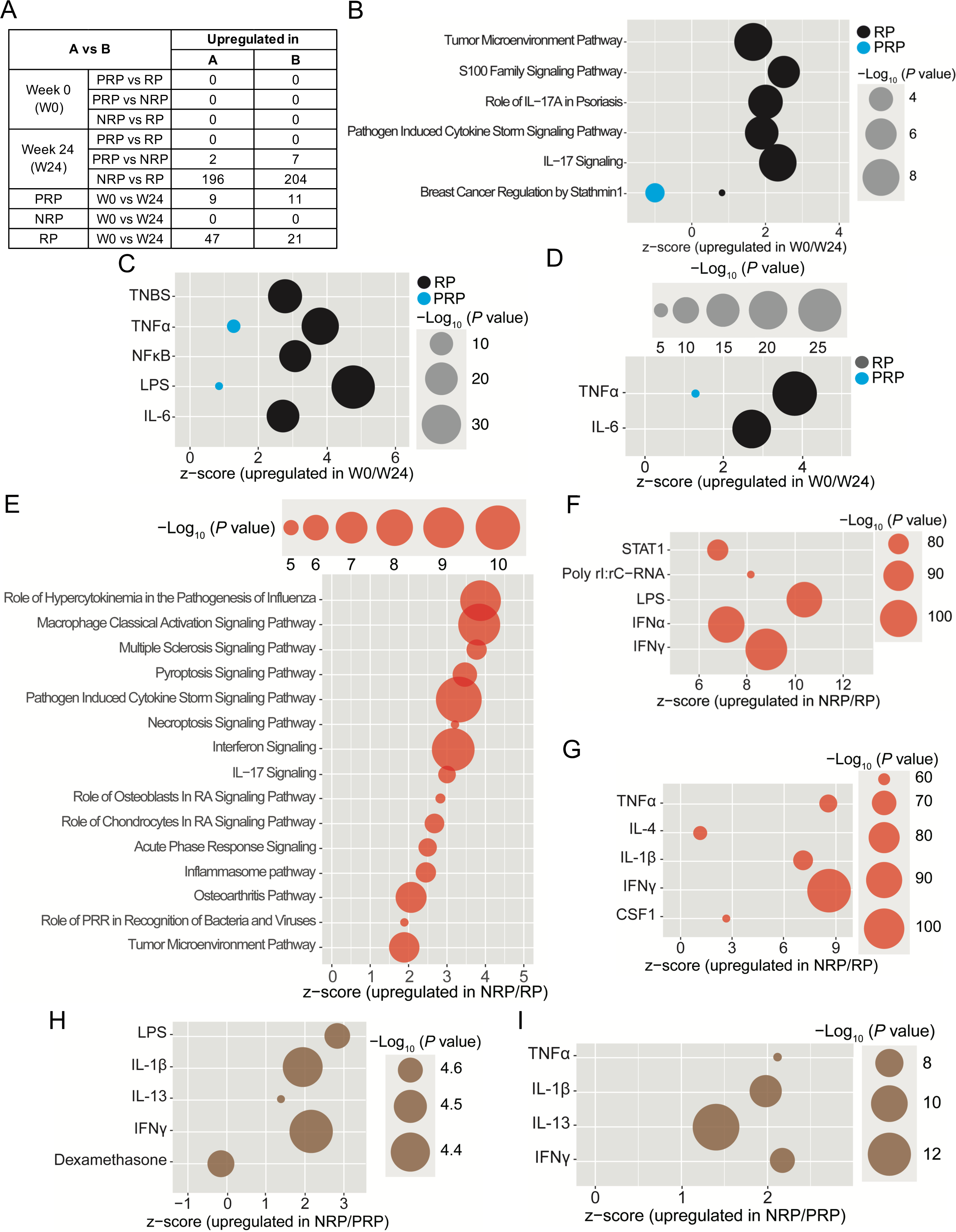
Tofacitinib treatment reduces inflammatory gene expression in responders. (A) Numbers of up- and down-regulated genes identified by transcriptome analysis in the indicated pair-wise comparison among responders (RPs), partial responders (PRPs), and nonresponders (NRPs) before [week (W)0] and after (W24) tofacitinib treatment. (B-D) Ingenuity pathway analysis (IPA) of the transcriptome of tofacitinib-receiving RP or PRP at W0 compared with W24. Top 6 pathways for the differentially expressed genes (DEGs) (average fold change > 2 and adjusted *P* < 0.05) in RP or PRP (B). Top 5 upstream regulators (C) and cytokines (D) related with inflammation activated in RP or PRP at W0 compared with W24. (E-I) IPA of the transcriptome of RP, PRP, or NRP at W24. Top 15 pathways for the DEGs (average fold change > 2 and adjusted *P* < 0.05) in NRP compared with RP. (E). Top 5 upstream regulator (F) and cytokines (G) related with inflammation activated in NRP compared with RP at W24. Top 5 upstream regulator (H) and cytokines (I) related with inflammation activated in NRP compared with PRP at W24. RP, responder; PRP, partial responder; NRP, nonresponder; W0, week 0; W24, week 24; RA, rheumatoid arthritis; PRR, pattern recognition receptor.

Pathway analyses of DEGs comparing W0 and W24 among RPs and PRPs included those associated with inflammation, such as cytokine signaling (Figure 1B-D). Genes related to the *Positive Regulation of Cytokine Production* (e.g., *NOS2*, *IL1B*, *TSLP*, and *IDO1*) were upregulated in RPs at W0 compared with W24 (Supplementary Figure 3D), confirming that clinical outcomes correlated with transcriptional changes in intestinal tissues of RPs. 20 genes were differentially expressed in PRPs at W0 compared with W24 (Supplementary Figure 3F), indicating that the effect of tofacitinib was less pronounced on the rectal transcriptome of PRPs.

At W24, immune signaling pathways such as *Interferon Signaling* were upregulated in NRPs compared with RPs (Figure 1E and Supplementary Figure 3E). Upstream regulators and cytokines such as STAT1 and IFNγ were more apparent in NRPs compared with RPs (Figure 1F and G). Inflammation-related upstream regulators and cytokines were similarly enriched in NRPs compared with PRPs, but the differences were less pronounced than when comparing RPs with NRPs (Figure 1H and I and Supplementary Figure 3G). Consistent with transcriptome analyses, an enzyme-linked immunosorbent assay (ELISA) showed that the concentration of fecal calprotectin, a biomarker for intestinal inflammation^46^, was decreased in RPs and PRPs at W24 compared with W0, whereas concentrations remained unchanged in NRPs (Supplementary Figure 3H).

Taken together, the above analyses reflect the clinical responsiveness of patients and validates their grouping into RP, PRP, and NRP. However, the baseline transcriptome does not distinguish patients in these categories.

### IBD-associated exome variants are common in RPs, PRPs, and NRPs

Exome sequencing identified 78 single nucleotide polymorphisms known to be associated with UC^47^ (Supplementary Table 4). Variants of the microbial sensor *NOD2* and transcription factors *FOXA1*, and *FOXA2* were enriched in UC patients who received tofacitinib compared with healthy controls (Supplementary Figure 4A). We further resolved exome variants by grouping into functional clusters according to gene ontology (GO) and observed those related to the epithelial barrier were associated with UC (Supplementary Figure 4B). We failed to find exome variants enriched in NRPs compared with RPs. Although these results do not rule out individual-specific roles for gene variants, they suggest that a mutation in the coding region of a gene is unlikely to underly tofacitinib responsiveness.

### Tofacitinib rescues viability of PDIOs stimulated with interferons

The above results indicate that the composition of the gut microbiome, rectal transcriptome, and exome variation do not predict tofacitinib responsiveness. As tofacitinib inhibits IFNγ or IFNβ-induced cell death of PDIOs^48–52^, we established colonic organoids derived from 23 UC patients at W0 in addition to 4 healthy donors (HDs) as negative controls (Supplementary Table 3). The viability and morphology of HD-derived organoids was unchanged up to 1 mM of tofacitinib treatment (Supplementary Figure 5A and B). Stimulation with IFNβ or IFNγ killed HD organoids, and tofacitinib rescued cell viability in a concentration-dependent manner (Supplementary Figure 5C and D). We reproduced these results using organoids derived from 6 UC patients, indicating that tofacitinib restores the number of viable HD and UC-derived organoids (Supplementary Figure 5E and F). A recent study demonstrated that removal of Noggin from the media sensitizes IL-17A-mediated cell death of human colonic organoids^23^. However, absence of growth factors Noggin and Wnt-3a did not impact the ability of tofacitinib to block IFNβ- or IFNγ-induced cell death of organoids (Supplementary Figure 6). Therefore, tofacitinib inhibits interferon-mediated cell death of HD- and UC-derived organoids.

### Tofacitinib-mediated inhibition of organoid cell death induced by TNFα and IFN predicts clinical responsiveness to tofacitinib

TNFα synergizes with IFNγ or IFNβ to induce JAK1/2- and STAT1-dependent death of intestinal organoids, potentially reproducing the microenvironment of an inflamed intestine^48, 49^. Thus, we tested the ability of tofacitinib to block cell death in organoids stimulated with TNFα+IFNβ or TNFα+IFNγ. The viability of both HD and UC organoids was rescued by 10 μM tofacitinib (Supplementary Figure 7A and B). In contrast, the degree to which 1 μM tofacitinib restored viability of organoids varied among UC organoids (Supplementary Figure 7B), indicating that a concentration of tofacitinib sufficient to inhibit cytokine-mediated death of control organoids is ineffective in a subset of UC organoids. Thus, we segregated UC organoids into groups based on their response to the drug. We refer to UC organoids in which 1 μM tofacitinib significantly increased viability in both co-stimulation conditions (TNFα+IFNβ and TNFα+IFNγ) as tofacitinib-sensitive (UC-S; Figure 2A and B). UC organoids insensitive to 1 μM tofacitinib (UC-IS) were divided into 2 subgroups: (1) organoids in which 1 μM tofacitinib failed to increase viability in both conditions (TNFα+IFNβ and TNFα+IFNγ) as UC-IS1, and (2) organoids in which 1 μM tofacitinib increased viability in the presence of TNFα+IFNβ, but not TNFα+IFNγ, as UC-IS2 (Figure 2C and D). There were no organoids in which tofacitinib increased viability in the presence of TNFα+IFNγ but not TNFα+IFNβ.

**Figure 2.**
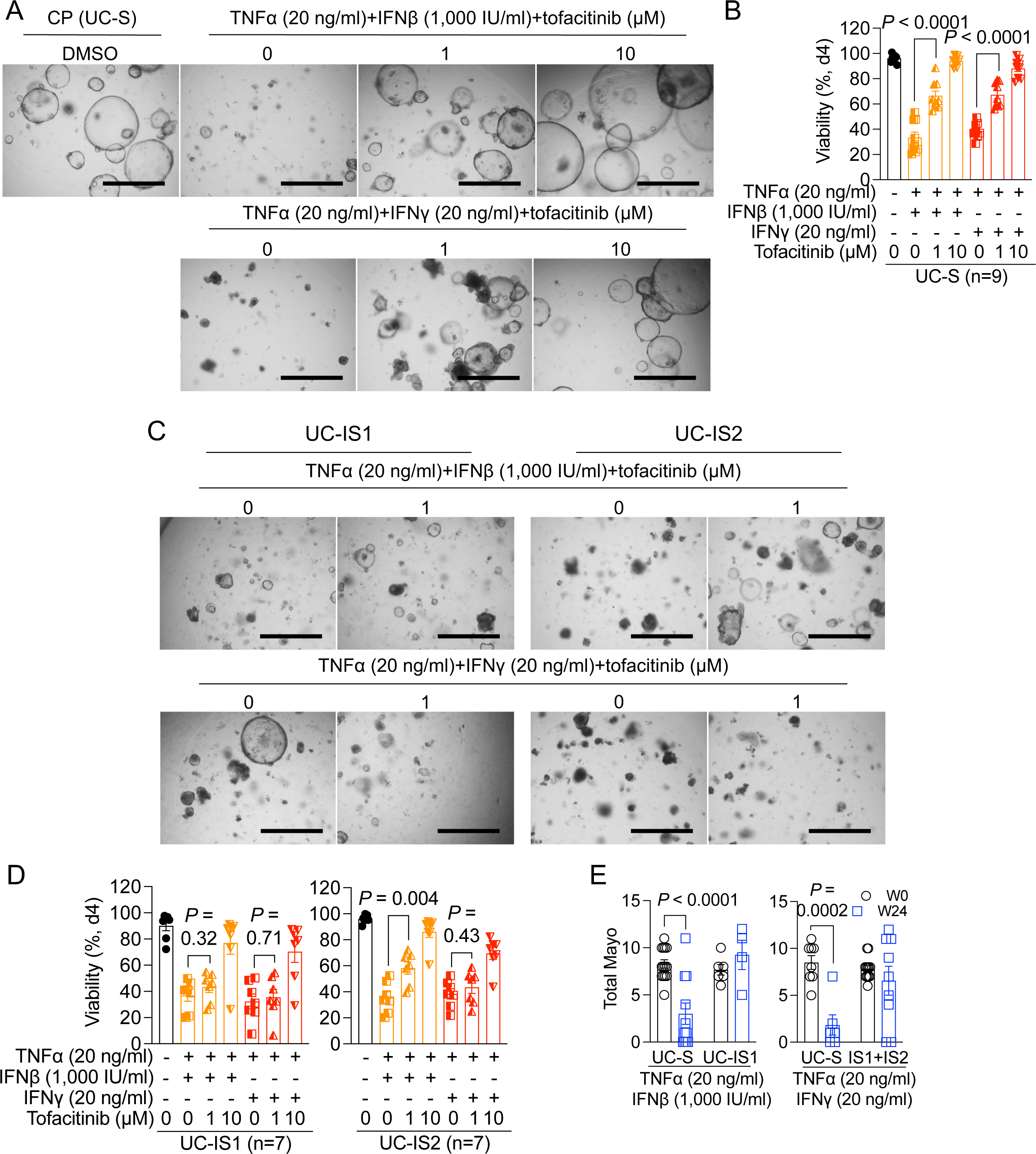
Patient-derived intestinal organoids display heterogenous sensitivity to tofacitinib. (A and B) Representative images (A) and quantification of viability (B) of tofacitinib-sensitive UC (UC-S) organoids stimulated with TNFα+IFNβ or TNFα+IFNγ in the presence of increasing concentration of tofacitinib. (C and D) Representative images (C) and quantification of viability (D) of group 1 (UC-IS1, left) or group 2 (UC-IS2, right) tofacitinib-insensitive UC organoids stimulated with TNFα+IFNβ or TNFα+IFNγ in the absence or presence of tofacitinib. (E) Changes in total mayo score between W0 and W24 according to the tofacitinib sensitivity of organoids stimulated with TNFα+IFNβ (left) or TNFα+IFNγ (right). Bars in A and C, 1,000 μm. Data points in B and D represent individual organoid lines and are mean of three technical replicates. Data points in E represent individual patients. Bars represent mean±standard error of mean (SEM). UC, ulcerative colitis; CP, carrier protein; DMSO, dimethyl sulfoxide. UC-S, tofacitinib-sensitive UC patient; UC-IS1, group 1 tofacitinib-insensitive UC patient; UC-IS2, group 2 tofacitinib-insensitive UC patient. Indicated *P* values by unpaired *t* test in B, D, and E.

As expected, stimulation of organoids with TNFα+IFNβ or IFNγ induced expression of target genes involved in feedforward signal amplification such as *STAT1*, *TNFα*, *ISG15*, *IFNβ1*, and *IFNλ2/3* (Supplementary Figure 7C). Adding 1 μM of tofacitinib generally suppressed this transcriptional response in HD and UC-S organoids, whereas tofacitinib was less effective in inhibiting the transcriptional response in UC-IS1 and UC-IS2 organoids (Supplementary Figure 7C). Thus, failure to rescue colonic organoid viability with tofacitinib is associated with the lower efficacy of the drug at reducing the cytokine-induced transcriptional response.

We then examined how the efficacy of tofacitinib to improve organoid viability related to clinical outcomes. We first compared patient responsiveness between TNFα+IFNβ treated organoids that were rescued by tofacitinib versus those that were insensitive (UC-S versus UC-IS1). Tofacitinib decreased the total Mayo score between W0 and W24 in patients corresponding to the UC-S organoids but not US-IS1 (Figure 2E). Next, we performed a similar comparison for organoids treated with TNFα+IFNγ. For this analysis, we grouped UC-IS1 and UC-IS2 because tofacitinib failed to rescue viability in both organoid subsets under this condition. Again, the total Mayo score was reduced in patients corresponding to the UC-S organoids but not US-IS organoids (Figure 2E). Therefore, the sensitivity of organoids to tofacitinib reflected individual clinical tofacitinib responsiveness of corresponding patients.

### STAT1 remains phosphorylated in tofacitinib-insensitive organoids

Tofacitinib inhibits STAT1 phosphorylation (pSTAT1)^53^, which we quantified by western blot. Cytokine stimulation induced pSTAT1 whereas total STAT1 was comparable across conditions (Figure 3A and B and Supplementary Figure 8). Treatment with 1 μM tofacitinib inhibited pSTAT1 in HD and UC-S organoids, but not in the UC-IS1 organoids (Figure 3A and B). Consistent with the partial viability rescue of UC-IS2 by tofacitinib, UC-IS2 organoids displayed decreased pSTAT1 induced by TNFα+IFNβ but not TNFα+IFNγ (Figure 3A and B). These results suggest that cell death occurs in cytokine-stimulated UC-IS organoids because 1 μM tofacitinib fails to reduce pSTAT1.

**Figure 3.**
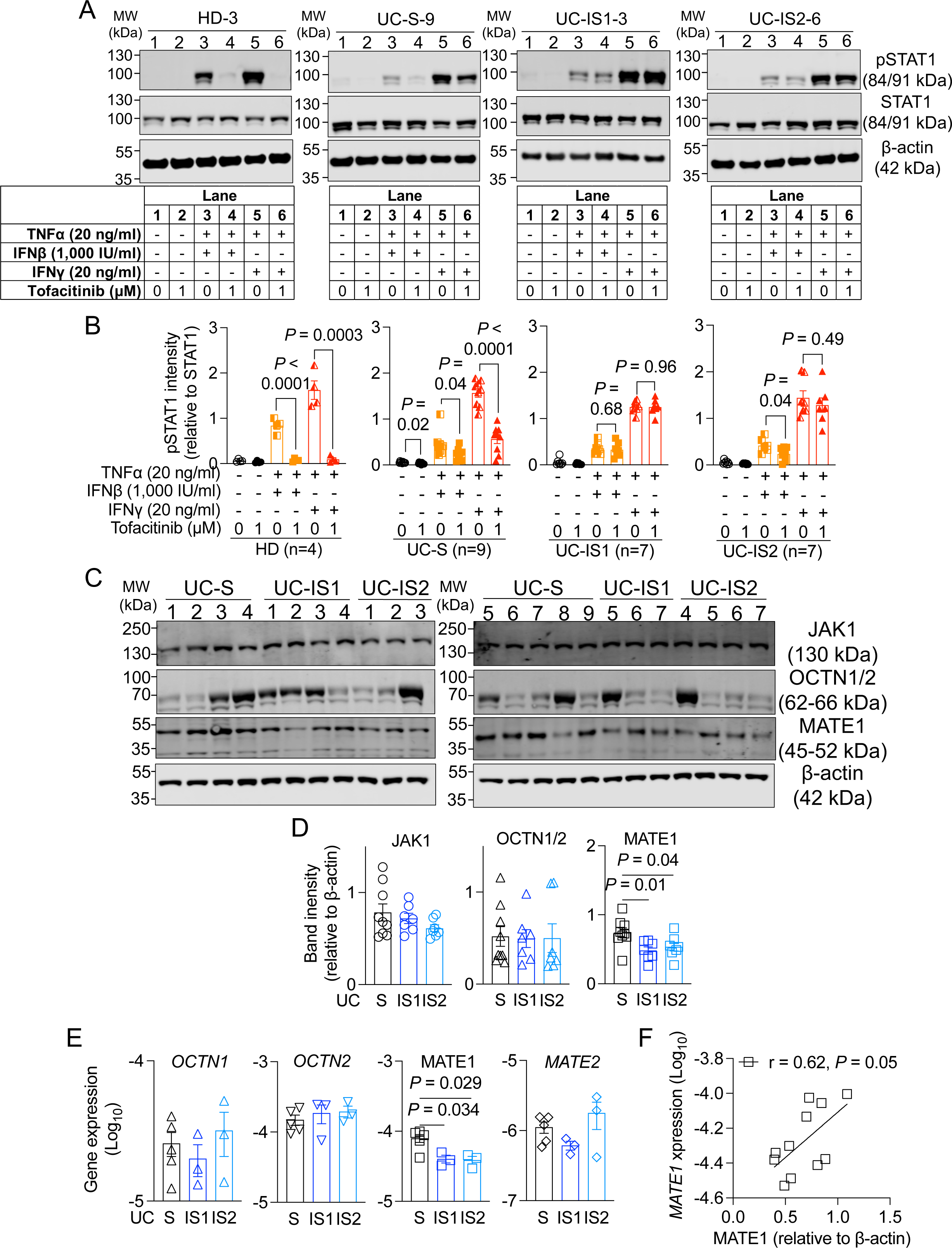
Insensitive organoids display uncontrolled STAT1 phosphorylation (pSTAT1) and low MATE1 expression. (A) Representative images of pSTAT1, STAT1, and β-actin in representative healthy donor (HD), UC-S, UC-IS1, or US-IS2 organoids stimulated with TNFα+IFNβ or TNFα+IFNγ in the presence of tofacitinib by western blot. The bottom tables represent treatment conditions of each lane. (B) pSTAT1 intensity relative to STAT1 in HD, UC-S, UC-IS1, or UC-IS2 organoids (from left to right). (C and D) Representative images of JAK1, OCTN1/2, MATE1, and β-actin (C) and quantification of band intensities of JAK, OCTN1/2, and MATE1 normalized to β-actin (D) in UC-S, UC-IS1, and UC-IS2 organoids treated with CP and DMSO by western blot. (E) RT-PCR analysis of *OCTN1*, *OCTN2*, *MATE1*, and *MATE2* expression in UC-S (n = 5), UC-IS1 (n = 3), and UC-IS2 (n = 3) organoids treated with mock CP and DMSO. (F) Correlation between *MATE1* expression and MATE1 intensity. Data points in B and D-F represent individual organoid lines. Data points in E are mean of two technical replicates. Bars represent mean±SEM. r, Pearson correlation coefficient. Indicated *P* values by unpaired *t* test in B, D, and E and simple regression analysis in F.

### Organoid sensitivity to tofacitinib is associated with increased levels of MATE1

A recent study demonstrated that a cationic transporter, MATE1, mediates uptake of tofacitinib^54^. Thus, we measured the level of JAK1 and MATE1 in unstimulated organoids by western blot. We also measured another cationic transporter, organic cation transporter novel (OCTN)1/2, as a control. UC-IS organoids displayed lower MATE1 than UC-S organoids while OCTN1/2 and JAK1 levels were comparable (Figure 3C and D). Transcript analysis reproduced western blot results: decreased *MATE1* expression in UC-IS organoids but comparable *OCTN1*, *OCTN2*, and *MATE2* expressions in UC-S and -IS organoids (Figure 3E). *MATE1* expression correlated with the protein level, indicating that differences in MATE1 between organoids occurs at the transcript level (Figure 3F). The transcript and protein levels of MATE1 were unchanged in organoids treated with cytokines with and without tofacitinib (Supplementary Figure 9). Thus, a decrease in MATE1 is associated with tofacitinib insensitivity in organoids.

We further tested whether MATE1-selective inhibitors^55^ prevent tofacitinib from rescuing the viability of UC-S organoids. We added MATE1-selective inhibitors (ritonavir, famotidine, or famotidine) or an OCT2-seletive inhibitor (pantoprazole) as a negative control to UC-S organoids stimulated with TNFα+IFNβ (Supplementary Figure 10A). The inhibitors were not toxic to both unstimulated and cytokine-stimulated organoids (Supplementary Figure 10B and C). Treatment with ritonavir, indinavir, and famotidine inhibited tofacitinib-mediated rescue of organoid viability in a concentration-dependent manner whereas tofacitinib remained effective in pantoprazole-treated organoids (Figure 4A and B). The same results were reproduced when stimulated with TNFα + IFNγ (Supplementary Figure 11).

**Figure 4.**
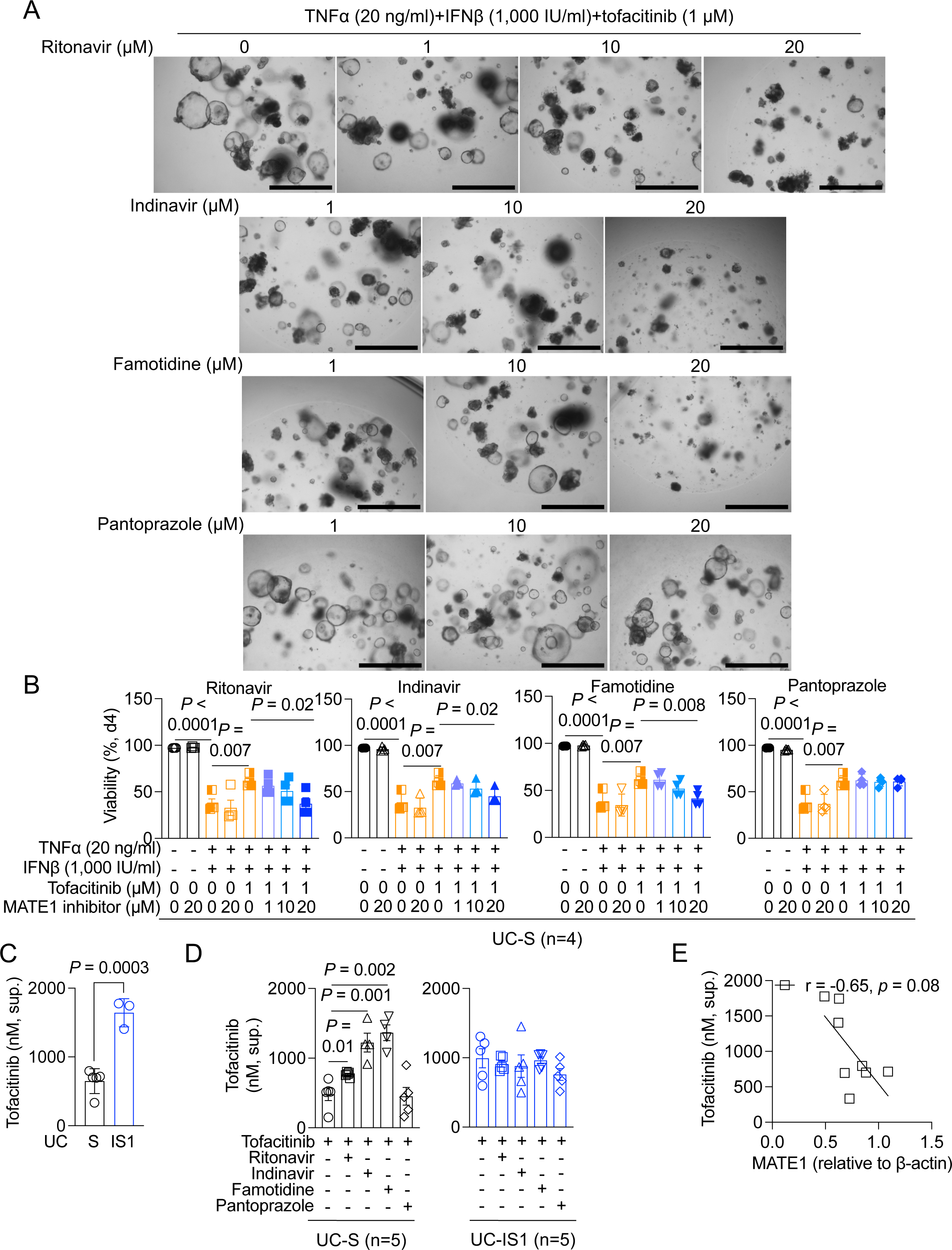
MATE1 mediates the uptake of tofacitinib in organoids. (A and B) Representative images (A) and quantification of viability (B) of UC-S organoids treated with TNFα+IFNγ+tofacitinib in the presence of increasing concentration of ritonavir, indinavir, famotidine, or pantoprazole. (C) Extracellular concentration of tofacitinib in UC-S or UC-IS1 organoids treated with 10 μM tofacitinib by LC/MS analysis. (D) Extracellular concentration of tofacitinib in UC-S or UC-IS1 organoids treated with 10 μM tofacitinib in the presence or absence of MATE1 inhibitors (20 μM) by LC/MS analysis. (E) Correlation between extracellular tofacitinib concentration and MATE1 expression in the organoids. Bars in A, 1,000 μm. Data points in B-E represent individual organoid lines. Bars represent mean±SEM. r, Pearson correlation coefficient. Indicated *P* values by unpaired *t* test in B-D and simple regression analysis in E.

To demonstrate that MATE1 is required for tofacitinib uptake, we measured remaining extracellular tofacitinib from organoids by liquid chromatography/mass spectrometer (LC/MS). Tofacitinib concentration in culture supernatants of UC-S organoids was significantly lower compared to UC-IS1 organoids (Figure 4C). The addition of MATE1 inhibitors increased the extracellular concentration of tofacitinib in UC-S but not in UC-IS1 organoids (Figure 4D). Lastly, the extracellular concentration of tofacitinib negatively correlated with MATE1 levels in organoids (Figure 4E). Thus, tofacitinib is imported into organoids in a MATE1-dependent manner, and this process is impaired in drug-insensitive organoids due to reduced MATE1 levels.

### MATE1 levels in organoids reflect the amount of MATE1 in primary tissues

To determine whether inter-individual differences in organoids reflect MATE1 levels in primary tissues, we measured MATE1 protein by immunofluorescence microscopy in colonic sections at W0 (baseline) from the same donors corresponding to individual organoid lines (Figure 5A and Supplementary Figure 12A-C). Consistent with a previous study^44^, MATE1 staining was scattered in the epithelium and lamina propria, but most intense along the apical brush border of epithelium (Figure 5A and Supplementary Figure 12A-C). Primary tissues displayed heterogeneous MATE1 protein levels (Figure 5B and Supplementary Figure 12). MATE1 was decreased in colonic sections from US-IS organoid donors compared with UC-S organoid donors (Figure 5C). Additionally, colonic sections of RPs displayed higher MATE1 levels than those of NRPs (Figure 5D). Lastly, MATE1 staining in colonic sections correlated with MATE1 intensity of organoids and was inversely associated with extracellular tofacitinib concentration of the organoids (Figure 5E). Because tissue specimens were procured prior to tofacitinib treatment, these results indicate that higher MATE1 levels in primary tissues were maintained in organoids and predict tofacitinib responsiveness.

**Figure 5.**
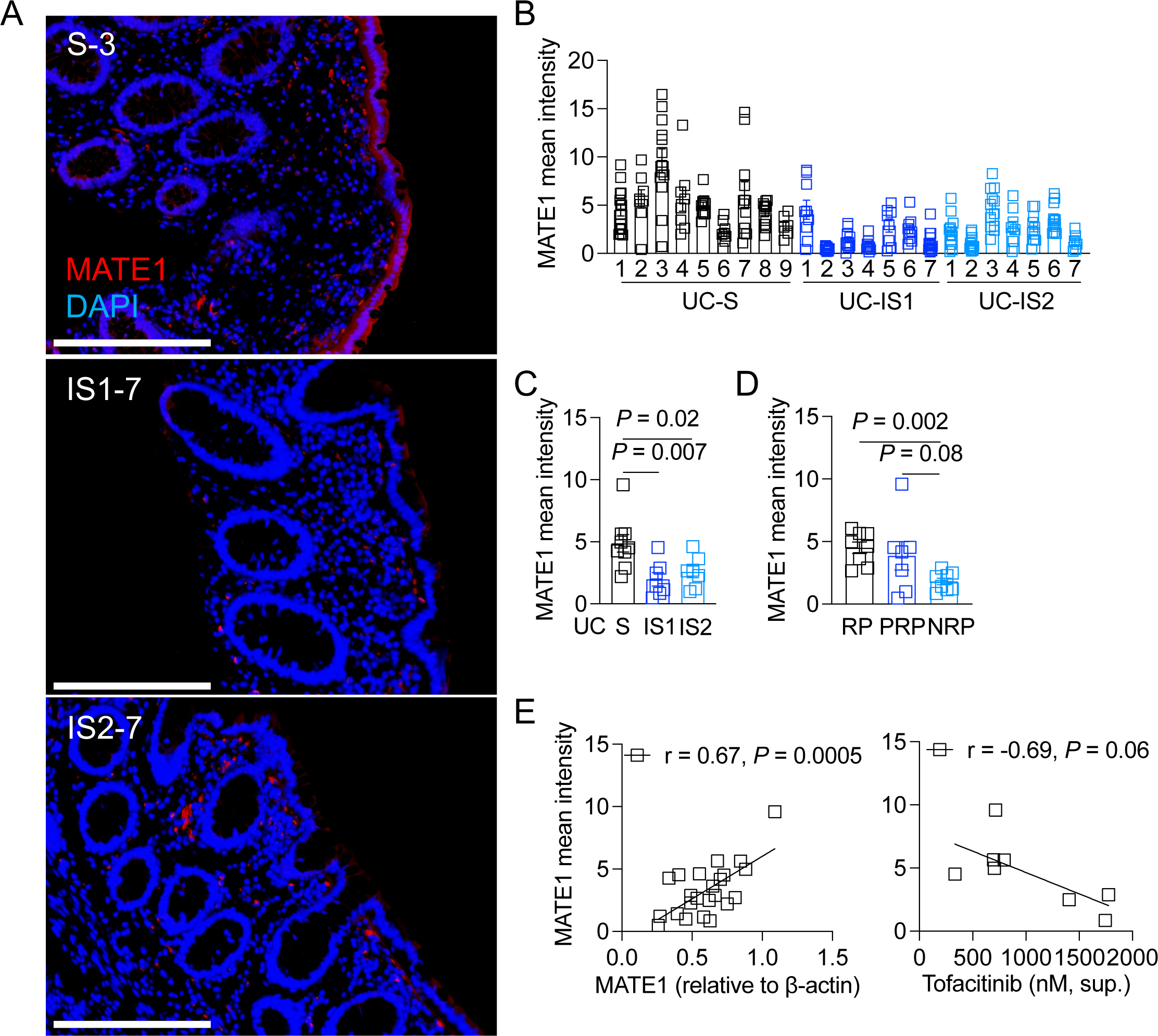
MATE1 expression in organoids reflect MATE1 levels in primary tissue and predict clinical outcome. (A) Representative images of MATE1 staining images in primary tissues of ascending colon form which UC-S-3 (upper), UC-IS1-7 (middle), UC-IS2-7 (lower) organoids were established. (B) Mean intensity of MATE1 per area in each field of view from primary tissues from which colonic organoids were established. (C and D) Mean intensity of MATE1 according to tofacitinib sensitivity in organoids (C) or tofacitinib responsiveness in patients (D). (E) Correlation between mean intensity of MATE1 in primary tissues and MATE1 intensity relative to β-actin in organoids (left) or extracellular concentration of tofacitinib (right). Bars in A, 200 μm. Data points in B are the field of views, and data points in C-E represent individual patients or organoids lines established from the same patients. Bars represent mean±SEM and at least two independent experiments were performed. r, Pearson correlation coefficient. Indicated *P* values by unpaired *t* test in C and D and simple regression analysis in E.

### JAK1 inhibitors upadacitinib and ruxolitinib rescue viability of tofacitinib-insensitive organoids

It is possible that other JAK inhibitors are efficacious in a setting in which tofacitinib is ineffective. Thus, we tested whether upadacitinib (JAK1 inhibitor), fedratinib (JAK2 inhibitor), or ruxolitinib (JAK1/2 inhibitor; Supplementary Figure 10D) rescued the cell death of UC-S and -IS organoids stimulated with rTNFα+rIFNβ. Using stored frozen organoids, we included tofacitinib as a control. We confirmed differences between UC-S and -IS1 organoids under tofacitinib treatment (Figure 6B and D). Upadacitinib and ruxolitinib increased the viability of UC-S and -IS organoids in a concentration-dependent manner whereas fedratinib exacerbated cytokine-induced cell death (Figure 6A-D). Similar results were obtained when stimulated with TNFα+IFNγ (Supplementary Figure 13).

**Figure 6.**
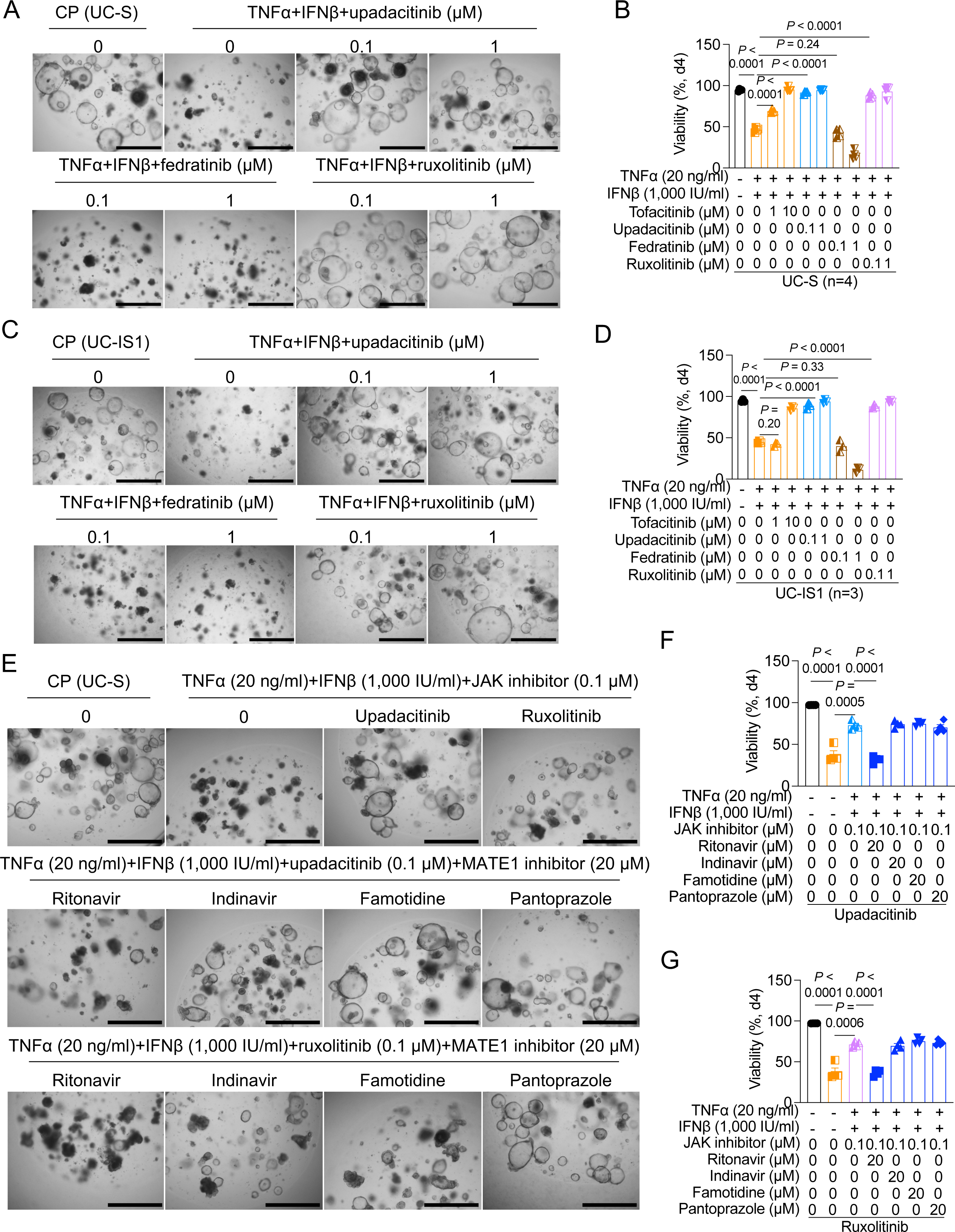
Upadacitinib and ruxolitinib rescue the cell death of UC-IS1. (A and B) Representative images (A) and quantification of viability (B) of UC-S organoids treated with TNFα+IFNβ in the presence of increasing concentration of tofacitinib, upadacitinib, or ruxolitinib on day 4. (C and D) Representative images (C) quantification of viability (D) of UC-IS1 organoids treated with TNFα+IFNβ in the presence of increasing concentration of tofacitinib, upadacitinib, or ruxolitinib on day 4. (E-G) Representative images (E) of quantification of viability (F and G) of upadacitinib- or ruxolitinib-treated US-S organoids stimulated with TNFα+IFNβ in the presence of ritonavir, indinavir, famotidine, or pantoprazole on day 4. Bars in A, C, and E, 1,000 μm. Concentration of cytokines in A and C are the same in B and D, respectively. Data points represent individual organoid lines and are mean of three technical replicates. Bars represent mean±SEM, and at least two independent experiments were performed. Indicated *P* values by unpaired *t* test in B, D, F, and G.

As MATE1 inhibitors also inhibit other transporters such as MATE2 and OCT1-3 with different half-maximal inhibitory concentration (IC_50_) values^55^, we tested whether MATE1 and OCT2 inhibitors eliminate the efficacy of updacitinib and ruxolitinib to rescue UC-S organoids. Of these inhibitors, ritonavir inhibited the activity of upadacitinib and ruxolitinib, resulting in failure to rescue the viability of organoids treated with TNFα+IFNβ and TNFα+IFNγ (Figure 6E-G and Supplementary Figure 14). Ritonavir inhibits OCT1 at lower concentrations than other inhibitors^55^, suggesting that uptake of upadacitinib and ruxolitinib may be dependent on OCT1. Thus, organoids insensitive to tofacitinib due to reduced MATE1 may respond to other JAK1 inhibitors.

## DISCUSSION

Prompt determination of drug responsiveness is a prerequisite for individualized drug selection, thereby improving precision medicine. In the present study, analyses of PDIOs and not the gut microbiota, rectal transcriptome, and exome variants of UC patients prior to treatment with tofacitinib predicted RPs and NRPs. The composition of the gut microbiota in RPs who achieved mucosal healing was unchanged during tofacitinib treatment, which is consistent with previous studies demonstrating that UC-associated dysbiosis in the microbiota is stable during short-term remission^56, 57^. In contrast, the rectal transcriptome at W24 differentiated RPs and NRPs, but baseline gene expression was similar across groups.

Recent findings documenting inter-individual differences between PDIOs led us to hypothesize that this system would demonstrate heterogeneous tofacitinib sensitivity, which we tested by examining cytokine-induced cell death. These experiments support a model in which tofacitinib uptake, dependent on *MATE1* expression, determines the clinical response to tofacitinib in patients with UC. Heterogeneity in *MATE1* expression was maintained in the primary tissues from which organoids were established. Our observations testing upadacitinib and ruxolitinib extend our model to other JAK inhibitors, showing that both drugs overcome the insensitivity of tofacitinib in PDIOs, our model highlighting the importance of transporter expression in determining IBD therapy responsiveness.

Transporters are determinants for drug disposition and response^58, 59^. MATE proteins of the SLC47 family are efflux transporters for clinically used drugs such as the antidiabetic agent metformin and the antibiotic cephalexin^60, 61^. Although MATEs (MATE1 and MATE2) are predominantly expressed in the human kidney, we demonstrated the expression of MATE1 in the colonic epithelium using PDIOs and their primary tissues. Our findings with selective MATE1 inhibitors may inform drug-drug interactions. Analyses of MATE1 inhibitors included famotidine (histamine-2 receptor antagonist) and pantoprazole (proton pump inhibitor), which are used to treat gastroesophageal reflux disease, a common comorbidity in patients with IBD^62^. As famotidine blocked the effect of tofacitinib by inhibiting MATE1-mediated uptake of tofacitinib (Figure 4 and Supplementary Figure 12), pantoprazole may be preferred during tofacitinib treatment^62–64^.

Previous studies have utilized PDIOs to predict drug responsiveness to anti-cancer and cystic fibrosis therapies ^28, 29, 65, 66^. Our findings with PDIOs from UC patients suggest that our approach may be extended to study mechanisms of drug response in other diseases and provide a foundation for discovering predictive biomarkers. In our study, heterogeneous *MATE1* expression in colonic organoids was retained in the primary tissues from which the organoids were generated (Figure 5 and Supplementary Figure 12), suggesting that MATE1 can be a biomarker for determining tofacitinib RPs. Quantification of biomarker intensity in primary tissues may be a straightforward option for predicting individual clinical efficacy of a given drug.

Although JAK inhibitors have diverse structures known to interact with multiple transporters^67, 68^, few studies have identified specific transporters involved in their uptake by relevant tissue types. The mechanism of transporter-dependent uptake of tofacitinib provides evidence that screening of transporter expression, and identification of specific transporters for uptake of JAK inhibitors, may be a tool for predicting individual-level treatment for multiple inflammatory diseases, including UC. A recent study demonstrated that baricitinib penetrates the cell membrane without an active transporter^54^. Thus, the combination of JAK inhibitors that target a broad spectrum of transporters or the development of transporter-independent JAK inhibitors may be an ideal therapeutic strategy that minimizes primary and secondary NRPs.

## Grant support

This work was supported in part by the Division of Intramural Research of NIAID/NIH (P’ng Loke), NIH grants K23DK124570 (Jordan Axelrad), P31CA016087, DK093668 (Ken Cadwell), AI121244 (Ken Cadwell), HL123340 (Ken Cadwell), AI130945 (Ken Cadwell), AI140754 (Ken Cadwell), and DK124336 (Ken Cadwell); Investigator-Initiated Research grant from Pfizer (Ken Cadwell and Jorden Axelrad); Faculty Scholar grant from the Howard Hughes Medical Institute (Ken Cadwell), Crohn’s & Colitis Foundation (Ken Cadwell and Jordan Axelrad), and Kenneth Rainin Foundation (Ken Cadwell). The funders had no role in study design, data collection and analysis, decision to publish, or preparation of the manuscript.

## Abbreviations

CFTR: (cystic fibrosis transmembrane conductance regulator)
DEG: (differentially expressed gene)
DMSO: (dimethyl sulfoxide)
EFG: (epidermal growth factor)
ELISA: (enzyme-linked immunosorbent assay)
FGF-2: (fibroblast growth factor 2)
HD: (healthy donor)
IC_50_: (half-maximal inhibitory concentration)
IF: (IGF-1 and FGF-2)
IFN: (interferon)
IGF-1: (insulin-like growth factor 1)
IPA: (ingenuity pathway analysis)
JAK: (Janus kinase)
LC/MS: (liquid chromatography/mass spectrometer)
MATE1: (multidrug and toxin extrusion protein 1)
NRP: (primary nonresponder)
NYU: (New York University)
OCTN: (organic cation transporter novel)
PCA: (principal component analysis)
PDIOs: (patient-derived intestinal organoids)
PRP: (partial responders)
PRR: (pattern recognition receptor)
pSTAT1: (STAT1 phosphorylation)
RA: (rheumatoid arthritis)
RP: (responder)
RPMI: (Roswell Park Memorial Institute)
RT: (real time)
SNP: (single nucleotide polymorphism)
STAT: (signal transducer and activator of transcription)
TNF: (anti-tumor necrosis factor)
UC-IS: (tofacitinib-insensitive UC)
UC-S: (tofacitinib-sensitive UC)
W: (week))

## Disclosures

Ken Cadwell has received research support from Pfizer, Takeda, Pacific Biosciences, Genentech, and Abbvie. Ken Cadwell has consulted for or received an honoraria from Puretech Health, Genentech, and Abbvie. Ken Cadwell is an inventor on U.S. patent 10,722,600 and provisional patent 62/935,035 and 63/157,225. Jordan E. Axelrad reports consultancy fees, honoraria, or advisory board fees from Abbvie, Adiso, Bristol Myers Squibb, Janssen, Pfizer, Fresnius, and BioFire Diagnostics.

## Acknowledgements

We would like to acknowledge NYU Grossman School of Medicine Reagent Preparation Core, Microscopy Laboratory, Genome Technology Center, and Center for Biospecimen Research and Development for use of their instruments and technical assistance. We also would like to thank Aravinda Chakravarti for his technical assistance of exome sequencing.

## Transcript Profiling and Writing Assistance

Not applicable in this study

## Authorship Contributions

Kyung Ku Jang, PhD (Conceptualization: Lead; Formal Analysis: Lead; Investigation: Lead; Methodology: Lead; Validation: Lead; Visualization: Lead; Writing – original draft: Lead; Writing – review & editing: Equal)

Defne Ercelen, BS (Formal Analysis: Equal; Investigation: Equal; Software: Lead; Visualization: Equal; Writing – original draft: Equal; Writing review & editing: Equal)

Jing Yu Carolina Cen Feng, BE (Formal Analysis: Equal; Resources: Equal; Validation: Equal; Writing – review & editing: Equal)

Sakteesh Gurunathan, MD (Formal Analysis: Equal; Methodology: Equal; Writing – original draft: Equal; Writing – review & editing: Equal)

Chaoting Zhou, BS (Formal analysis: Equal; Methodology: Equal; Writing – review & editing: Equal)

Aryeh Korman, BS (Formal Analysis: Equal; Methodology: Equal; Writing – original draft: Equal; Writing – review & editing: Equal)

Luke Newell, BS (Resources: Equal; Writing – review & editing: Equal).

David Hudesman, MD (Funding acquisition: Lead; Resources: Lead; Writing – review & editing: Equal)

Drew R. Jones, PhD (Methodology: Equal; Project administration: Equal; Writing – review & editing: Equal)

P’ng Loke, PhD (Funding acquisition: Equal; Writing – review & editing: Equal)

Jordan E. Axelrad, MD, MPH (Conceptualization: Lead; Funding acquisition: Lead; Project administration: Lead; Resources: Lead; Supervision: Lead; Writing – original draft: Lead; Writing – review & editing: Equal)

Ken Cadwell, PhD (Conceptualization: Lead; Funding acquisition: Lead; Investigation: Lead; Project administration: Lead; Resources: Lead; Supervision: Lead; Writing – original draft: Lead; Writing – review & editing: Equal)

## Data Transparency Statement

16S rRNA sequencing and exome sequencing data are deposited in National Center for Biotechnology Information Sequence Read Archive with accession numbers PRJNA1024576 and PRJNA1034220, respectively. Gene expression raw data are available from Gene Expression Omnibus with an accession number GSE245890. The clinical trial is preregistered in ClinicalTrial.gov with identifier NCT03663400. All data obtained and/or analyzed in this study are available from the corresponding authors upon reasonable request.

---

**Supplementary Figure 1.**
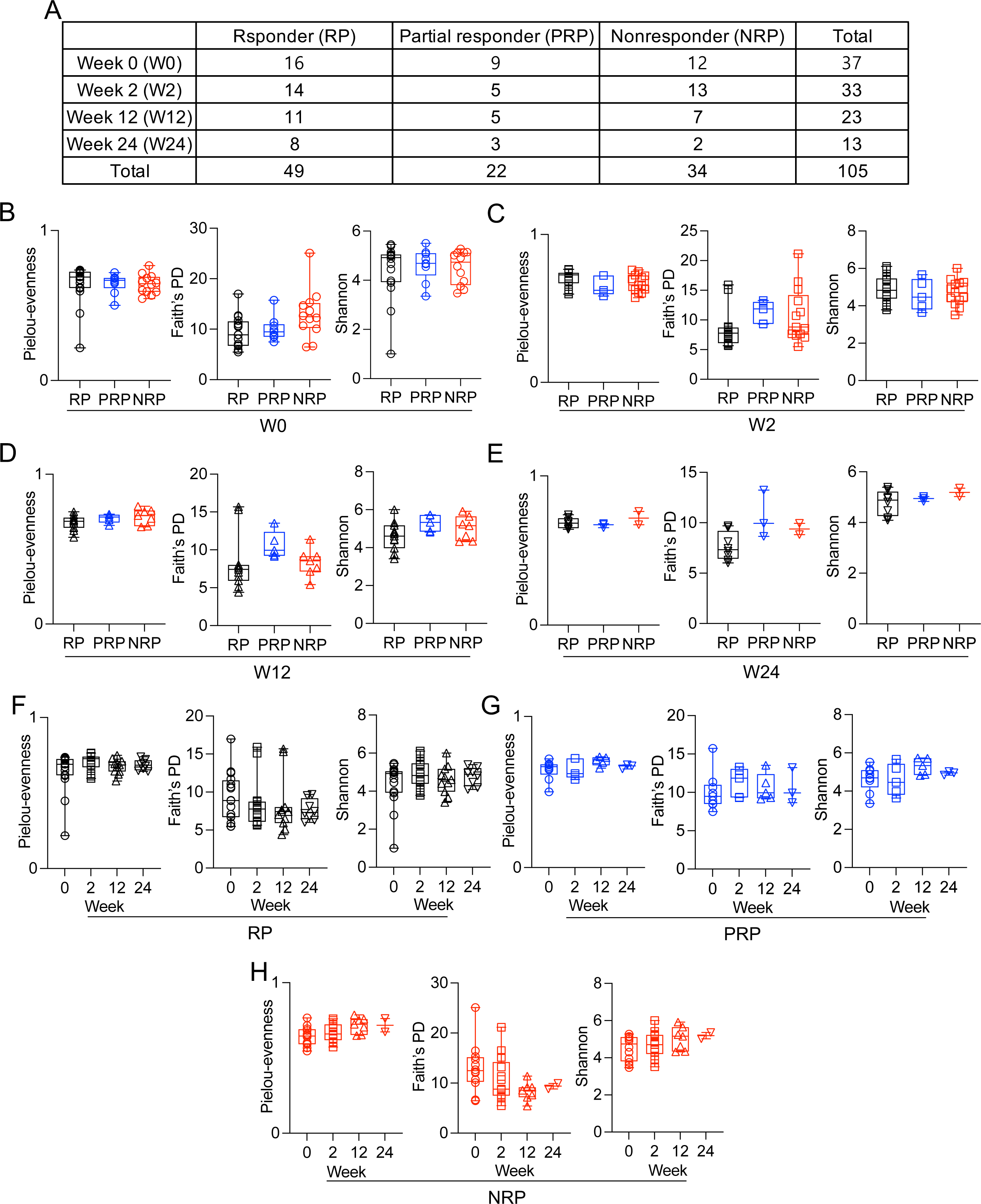
Microbiome analysis of stool samples from ulcerative colitis (UC) patients receiving tofacitinib. (A) Numbers of fecal samples collected from RPs, PRPs, and NRPs according to collection time points W0, 2, 12, and 24. (B-H) 16S rRNA sequencing of stool from UC patients in A. Alpha diversity values calculated as Pielou-evenness (left), Faith’s phylogenetic diversity (PD) (middle), and Shannon (right) indices of RP, PRP, and NRP on W0 (B), 2 (C), 12 (D), or 24 (E). Longitudinal changes in Pielou-evenness (left), Faith’s PD (middle), and Shannon (right) indices of RP (F), PRP (G), and NRP (H). Data points in B-H represent individual patients. RP, responder; PRP, partial responder; NRP, nonresponder; W0, week 0; W2, week 2; W12, week 12; W24, week 24.

**Supplementary Figure 2.**
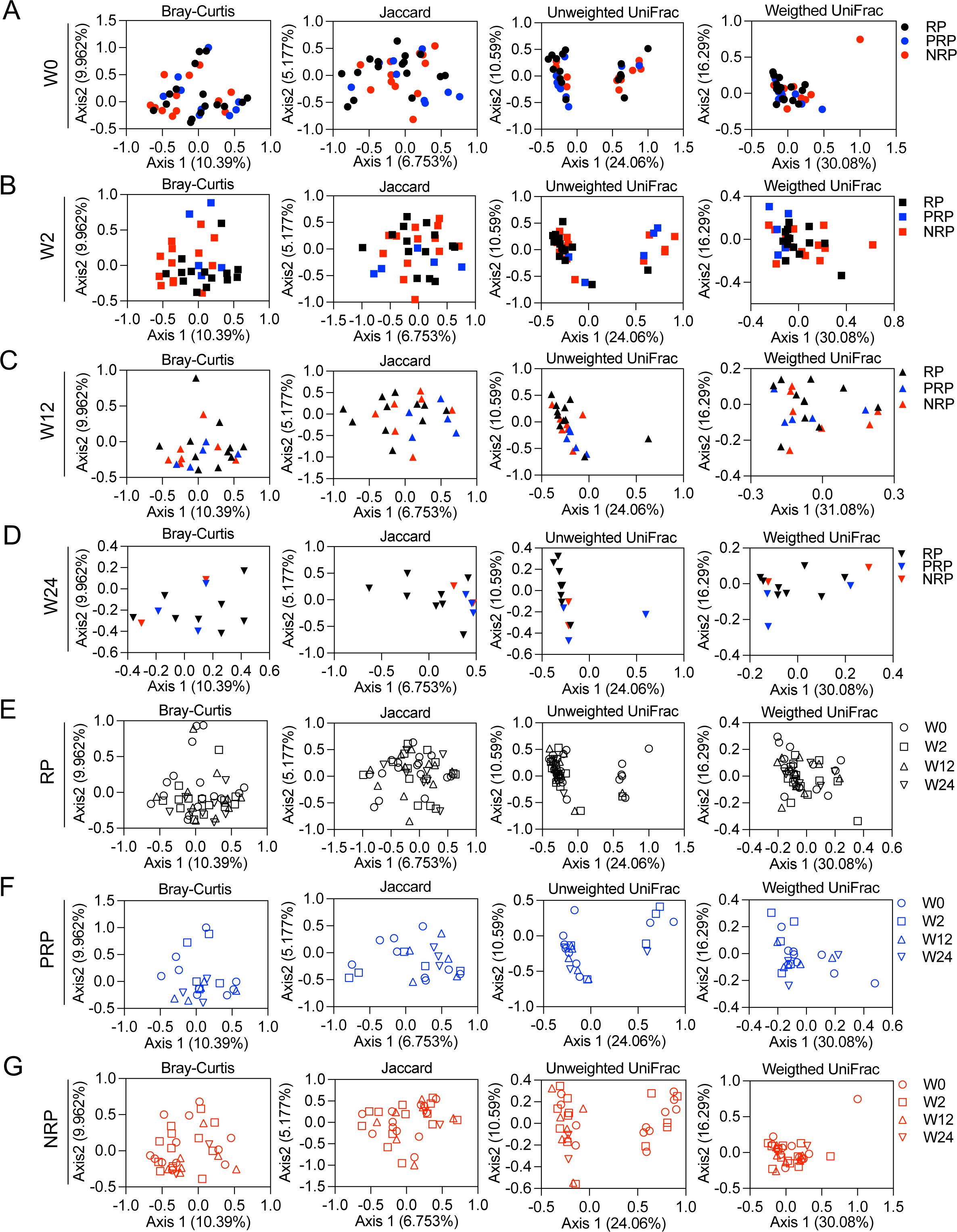
Beta diversity of microbiome from tofacitinib-treated UC patients. (A-G) 16S rRNA sequencing of stool from UC patients in Supplementary figure 1A. Principal coordinate analyses of beta diversity determined by Bray-Curtis, Jaccard, and Unweighted and Weighted UniFrac methods (from left to right). Beta diversity of RP, PRP, and NRP on W0 (A), W2 (B), W12 (C), or W24 (D). Longitudinal changes in beta diversity of RP (E), PRP (F), and NRP (G). Data points in A-G represent individual samples.

**Supplementary Figure 3.**
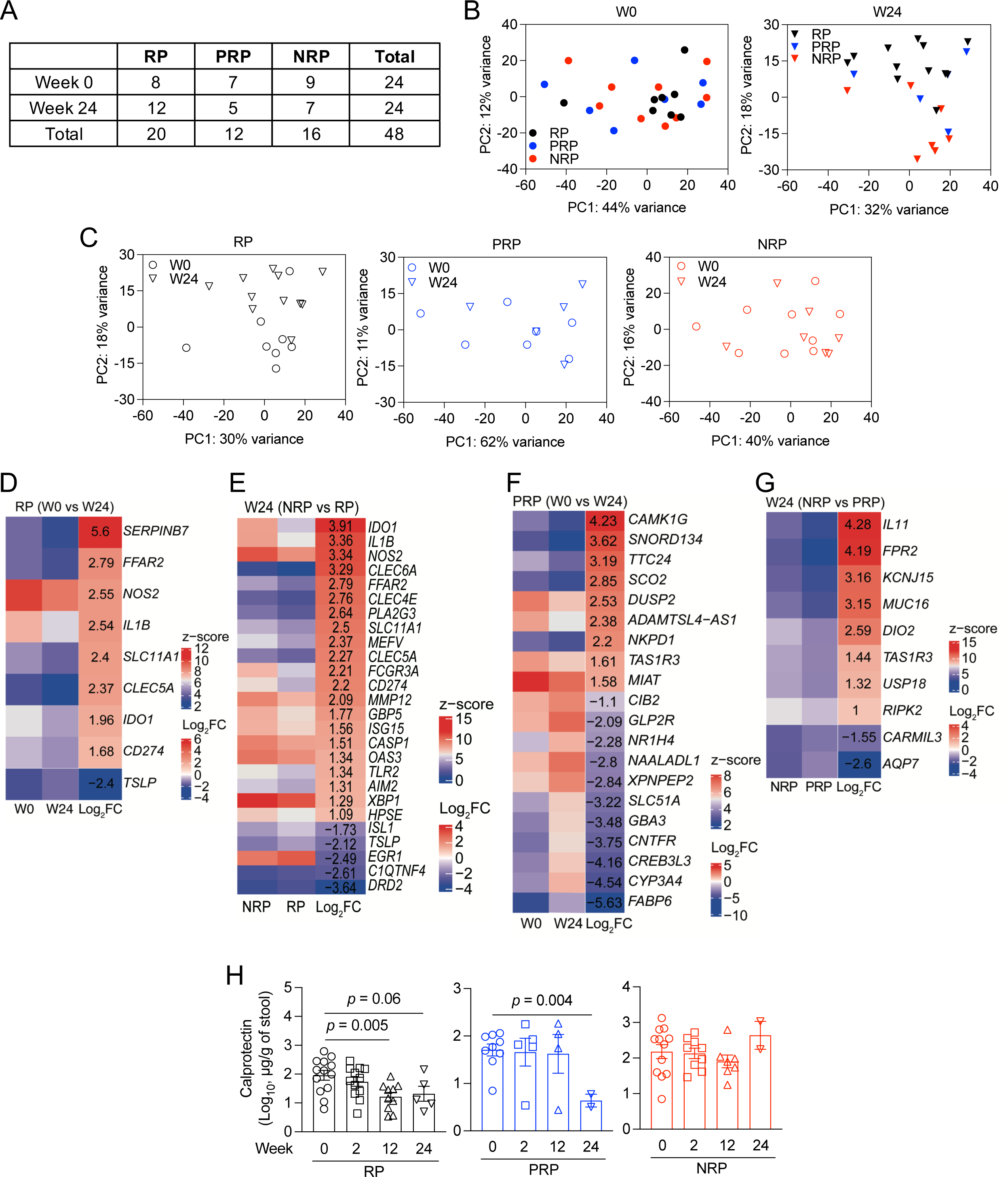
Analysis of rectal transcriptome and fecal calprotectin in tofacitinib-enrolled UC patients. (A) Numbers of rectal pinch biopsies collected from RP, PRP, and NRP at W0 (left) or W24 (right). (B) Unsupervised clustering based on expression of most variable genes by clinical responsiveness of tofacitinib (RP, PRP, and NRP) at W0 (left) or W24 (right). (C) Unsupervised clustering based on expression of most variable genes by collection time points of RP (left), PRP (middle), or NRP (right). (D and E) Heatmaps displaying normalized z-score and Log_2_ fold change (FC) of differentially expressed genes (DEGs) (average fold change > 2 and adjusted *P* value < 0.05) in RP at W24 compared with W0 (F) or RP compared with NRP at W24 annotated in gene ontology (GO): 0001819. (F and G) Heatmaps displaying normalized z-score and Log_2_ FC of DEGs (average fold change > 2 and adjusted *P* value < 0.05) in PRP at W24 compared with W0 (D) or in PRP compared with NRP at W24 (E). (H) Longitudinal changes in fecal calprotectin concertation of RP (left), PRP (middle), and NRP (right) in Supplementary figure 1A. Data points in B, C, and H represent individual patients and data points in H are mean of technical duplicates. Bars represent mean±SEM. Indicated *P* values by unpaired *t* test in H.

**Supplementary Figure 4.**
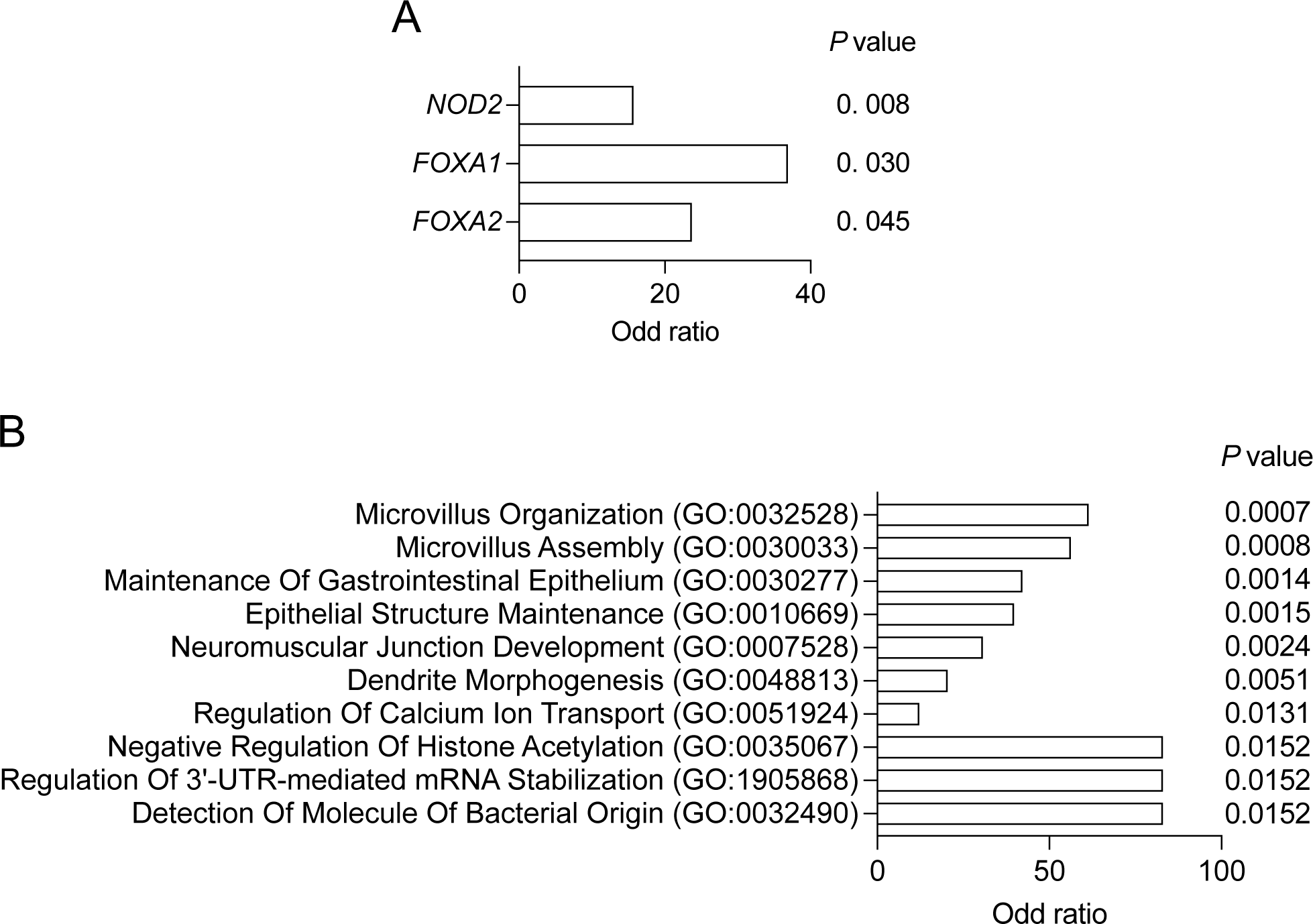
Analysis of exome variants in tofacitinib-enrolled UC patients. (A and B) Odds ratio for transcription factors (A) and top 10 GOs that significantly contribute to the total variance explained in IBD. *P* values are indicated on the right of each panel.

**Supplementary Figure 5.**
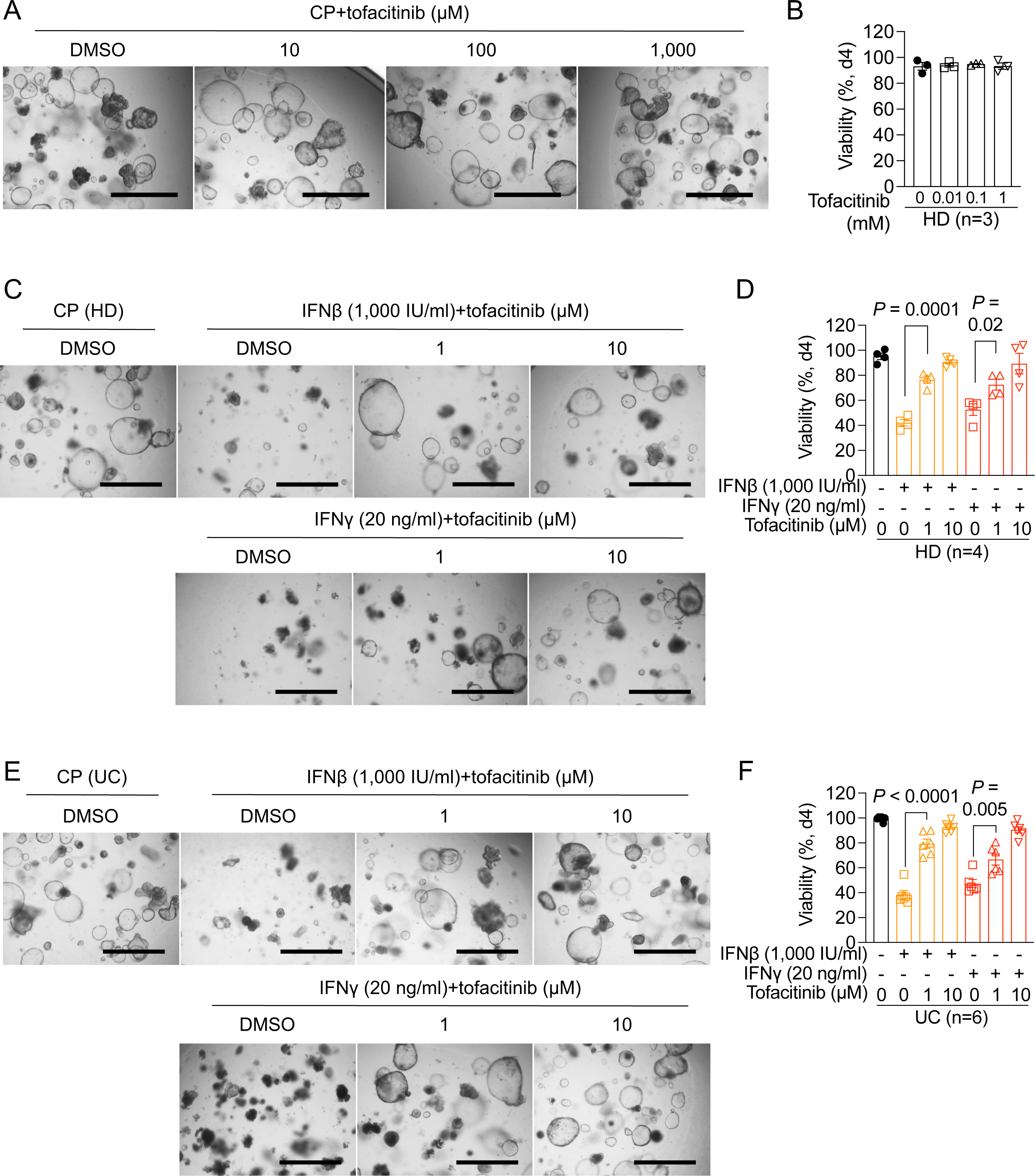
Effect of tofacitinib on viability of unstimulated or interferon-treated organoids. (A and B) Representative images (A) and quantification of the proportion of viable HD organoids by microscopy (B) in the presence of increasing concentration of tofacitinib on day 4. (C and D) Representative images (C) and quantification of viability (D) of HD organoids stimulated with either IFNβ or IFNγ in the presence of increasing concentration of tofacitinib on day 4. (E and F) Representative images (E) and quantification of viability (F) of UC organoids stimulated with either IFNβ or IFNγ in the presence of increasing concentration of tofacitinib on day 4. Bars in A, C, and E, 1,000 μm. Concentrations of cytokines and tofacitinib are indicated in each panel. Data points in B, D, and F represent individual organoid lines and are mean of three technical replicates. Bars represent mean±SEM. HD, healthy donor; UC, ulcerative colitis; CP, carrier protein; DMSO, dimethyl sulfoxide. Indicated *P* values by unpaired *t* test in D and F.

**Supplementary Figure 6.**
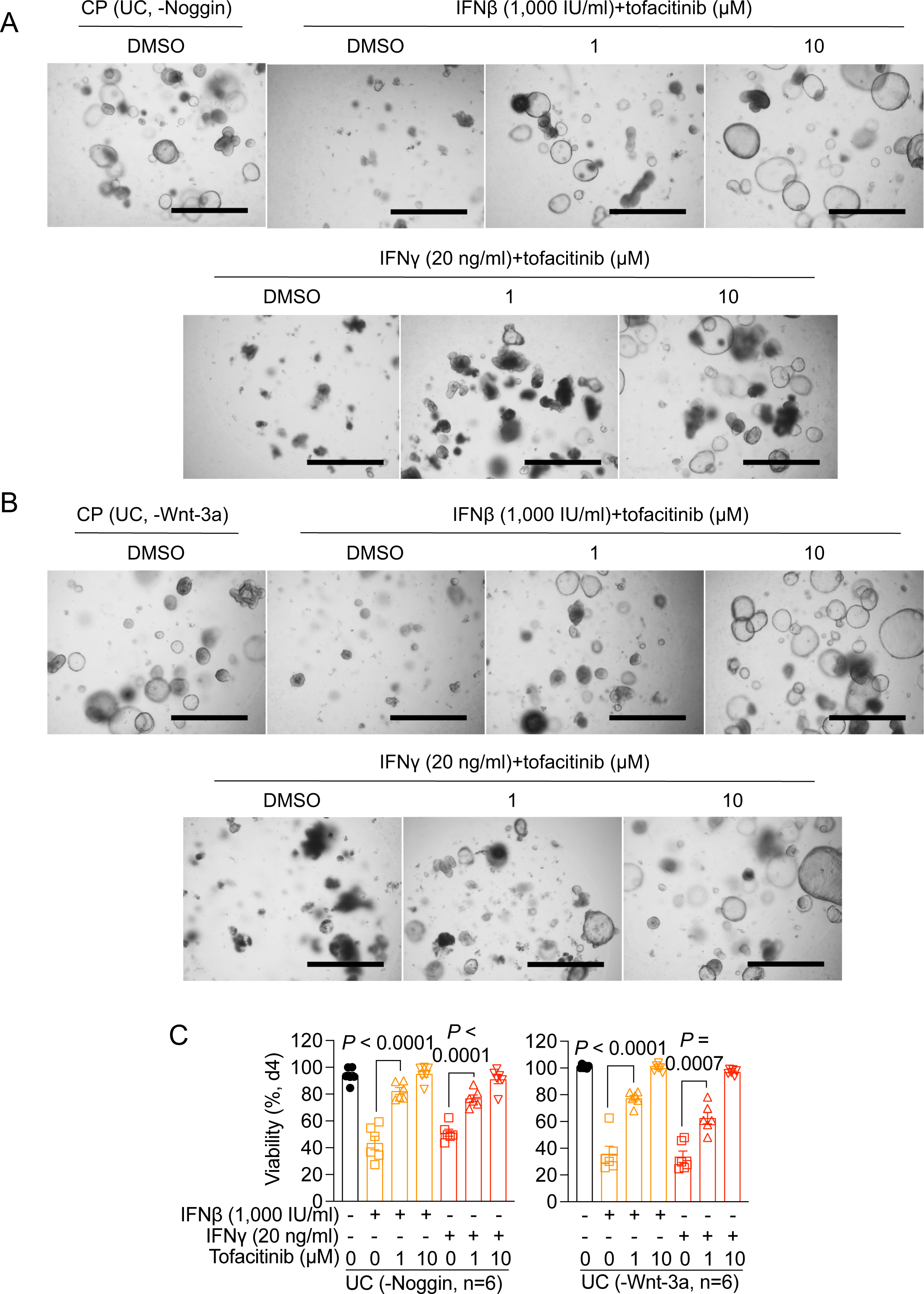
Tofacitinib-mediated increases in organoid viability are retained in in the absence of Noggin and Wnt-3a. (A and B) Representative images of UC organoids stimulated with either IFNβ or IFNγ in the absence of Noggin (A) or Wnt-3a (B). (C) Quantification of viability of organoids in A and B in the absence of either Noggin (left) or Wnt-3a (right). Bar in A and B, 1,000 μm. Concentrations of cytokines and tofacitinib are indicated in each panel. Data points in C represent individual organoid lines and are mean of three technical replicates. Bars represent mean±SEM. Indicated *P* values by unpaired *t* test in C

**Supplementary Figure 7.**
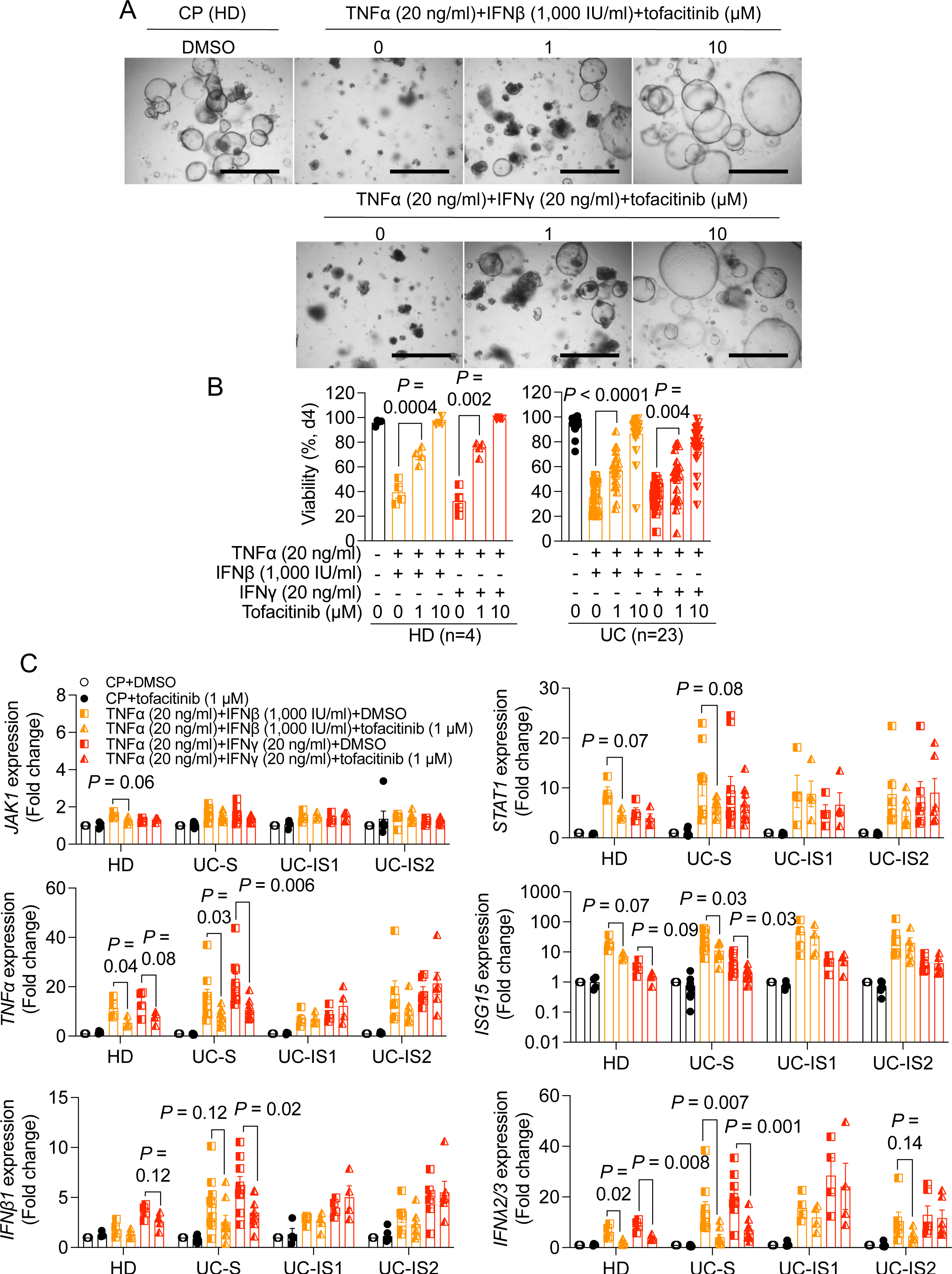
Analyses of viability and transcripts of HD and UC patient-derived organoids treated with cytokines and tofacitinib. (A) Representative images of HD organoids stimulated with TNFα+IFNβ or TNFα+IFNγ in the presence of increasing concentration of tofacitinib on day 4. (B) Quantification of viability from HD (left) or UC (right) organoids stimulated with TNFα+IFNβ or TNFα+IFNγ in the presence of increasing concentration of tofacitinib on day 4. (C) qRT-PCR analysis of *JAK1*, *STAT1*, *TNFα*, *ISG15*, *IFNβ1*, and *IFNλ2/3* in HD organoids (n = 4) or UC-S (n = 9), UC-IS1 (n = 4), and UC-IS2 (n = 6) organoids stimulated with TNFα+IFNβ or TNFα+IFNγ in the presence or absence of 1 μM tofacitinib for 4 hr. Bar in A, 1,000 μm. Concentrations of cytokines and tofacitinib are indicated in each panel. Data points in A and C represent individual organoid lines and are mean of at least two technical replicates. Bars represent mean±SEM. UC-S, tofacitinib-sensitive UC patient; UC-IS1, group 1 tofacitinib-insensitive UC patient; UC-IS2, group 2 tofacitinib-insensitive UC patient. Indicated *P* values by unpaired *t* test in B and C.

**Supplementary Figure 8.**
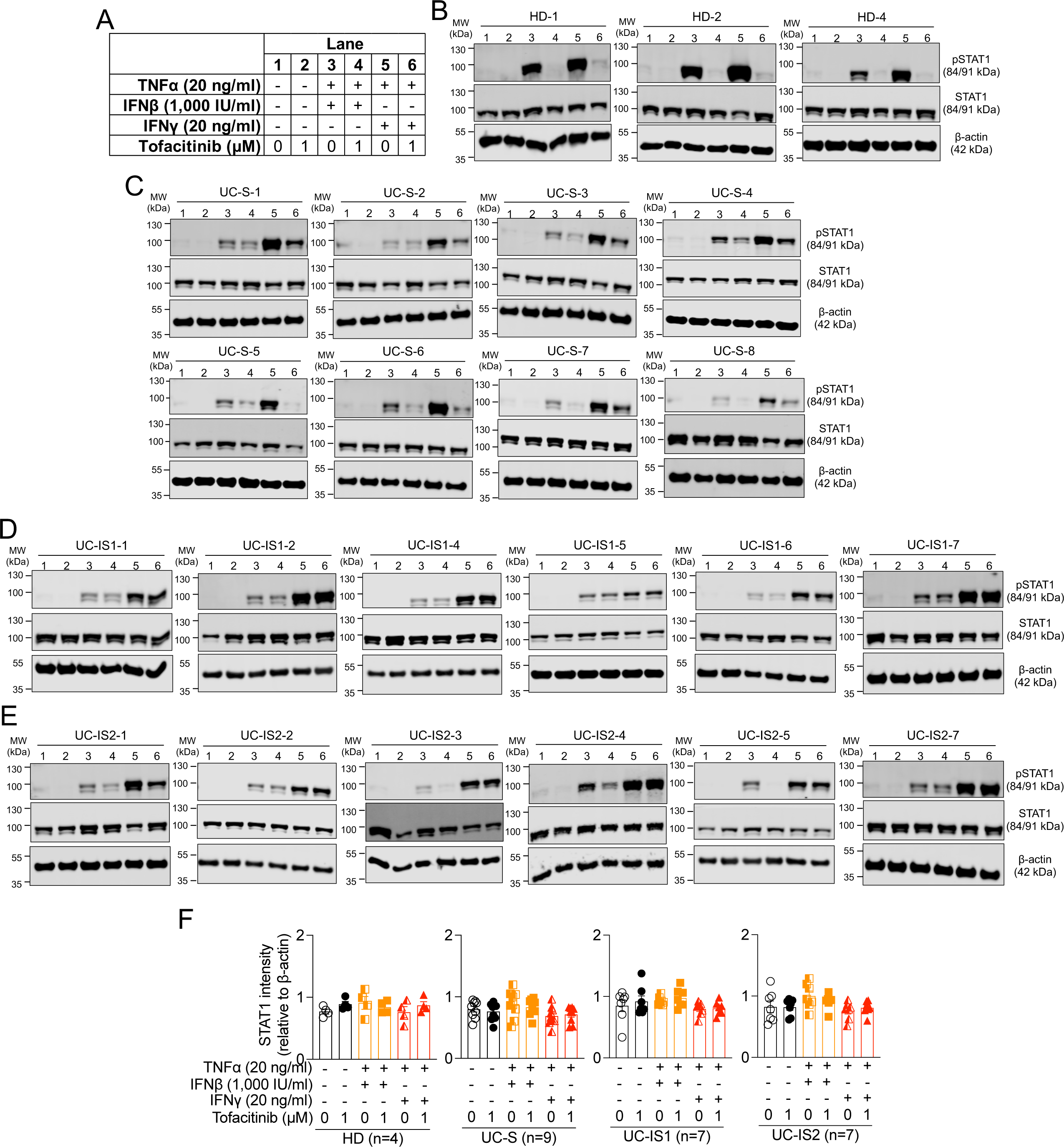
Western blot analysis of UC patient-derived organoids. (A) Table represents treatment conditions in B-E. (B-E) Representative images of pSTAT1, STAT1, and β-actin in HD (HD-1, 2, and 4, A), UC-S (S-1 to 8, B), UC-IS1 (IS1-1, 2, and 4 to 7, C), or US-IS2 (IS2-1 to 5 and 7, D) organoids stimulated with TNFα+IFNβ or TNFα+IFNγ in the presence or absence of tofacitinib for 4 hr by western blot. Images of remaining organoid lines are included in Figure 3A. (F) STAT1 intensity relative to β-actin in HD, UC-S, UC-IS1, or UC-IS2 organoids (from left to right). Data points in F represent individual organoid lines. Bars represent mean±SEM.

**Supplementary Figure 9.**
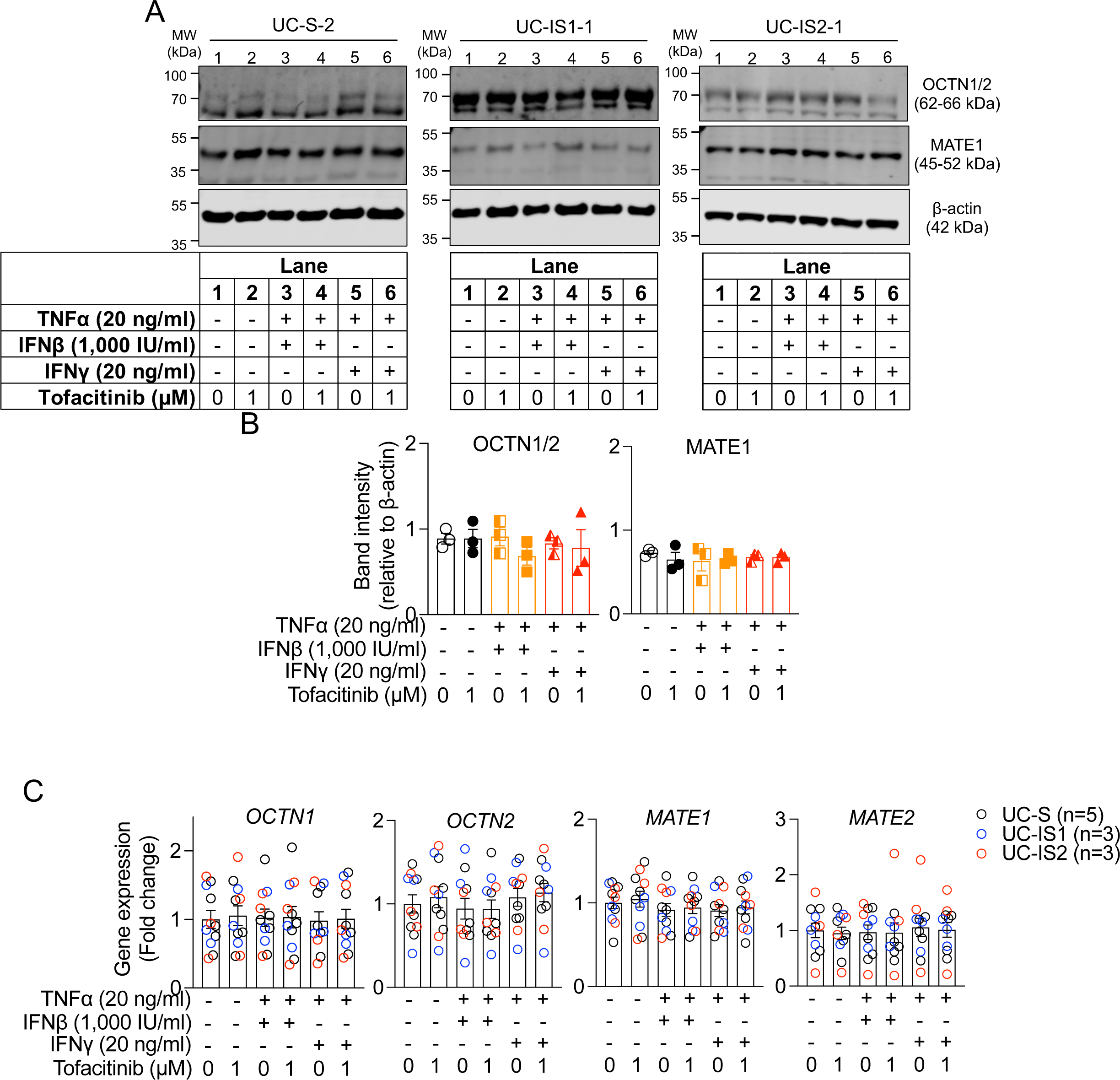
Cytokine stimulation and tofacitinib treatment does not affect MATE1 levels. (A) Representative images of OCTN1/2, MATE1, and β-actin in UC-S-2 (left), UC-IS1-1 (middle), or US-IS2-1 (right) organoids stimulated with TNFα+IFNβ or TNFα+IFNγ in the presence or absence of tofacitinib for 4 hr by Western blot. The bottom tables represent treatment conditions of each lane. (B) OCTN1 (left) or MATE1 (right) intensity relative to β-actin in UC-S, UC-IS1, or UC-IS2 organoids in B.(C) RT-PCR analysis of *OCTN1*, *OCTN2*, *MATE1*, and *MATE2* in UC-S, UC-IS1, or UC-IS2 organoids stimulated with TNFα+IFNβ or TNFα+IFNγ in the presence or absence of tofacitinib for 4 hr. Concentrations of cytokines and tofacitinib are indicated in each panel. Data points in B and C represent individual organoid lines. Bars represent mean±SEM.

**Supplementary Figure 10.**
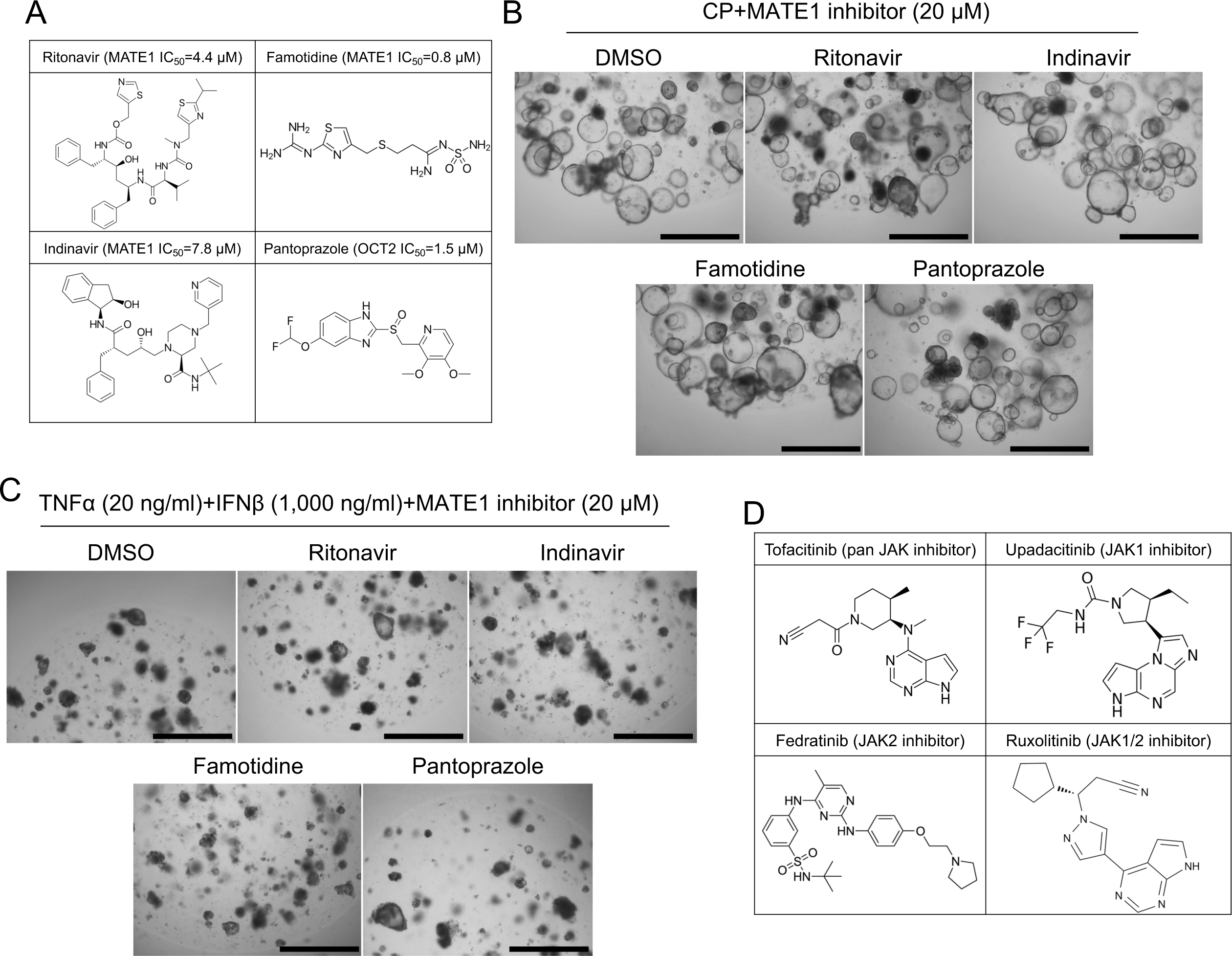
MATE1 inhibitors does not affect the viability of unstimulated organoids and structure of MATE1 inhibitors and JAK inhibitors. (A) Structure and half-maximal inhibitory concentration (IC_50_) of MATE1 inhibitors. (B and C) Representative images of unstimulated (B) or TNFα and IFNβ-stimulated (C) organoids in the presence of MATE1 inhibitors on day 4. (C) Structure and targets of JAK inhibitors. Concentrations of cytokines and MATE1 inhibitors are indicated in B and C. IC_50_, half-maximal inhibitory concentration.

**Supplementary Figure 11.**
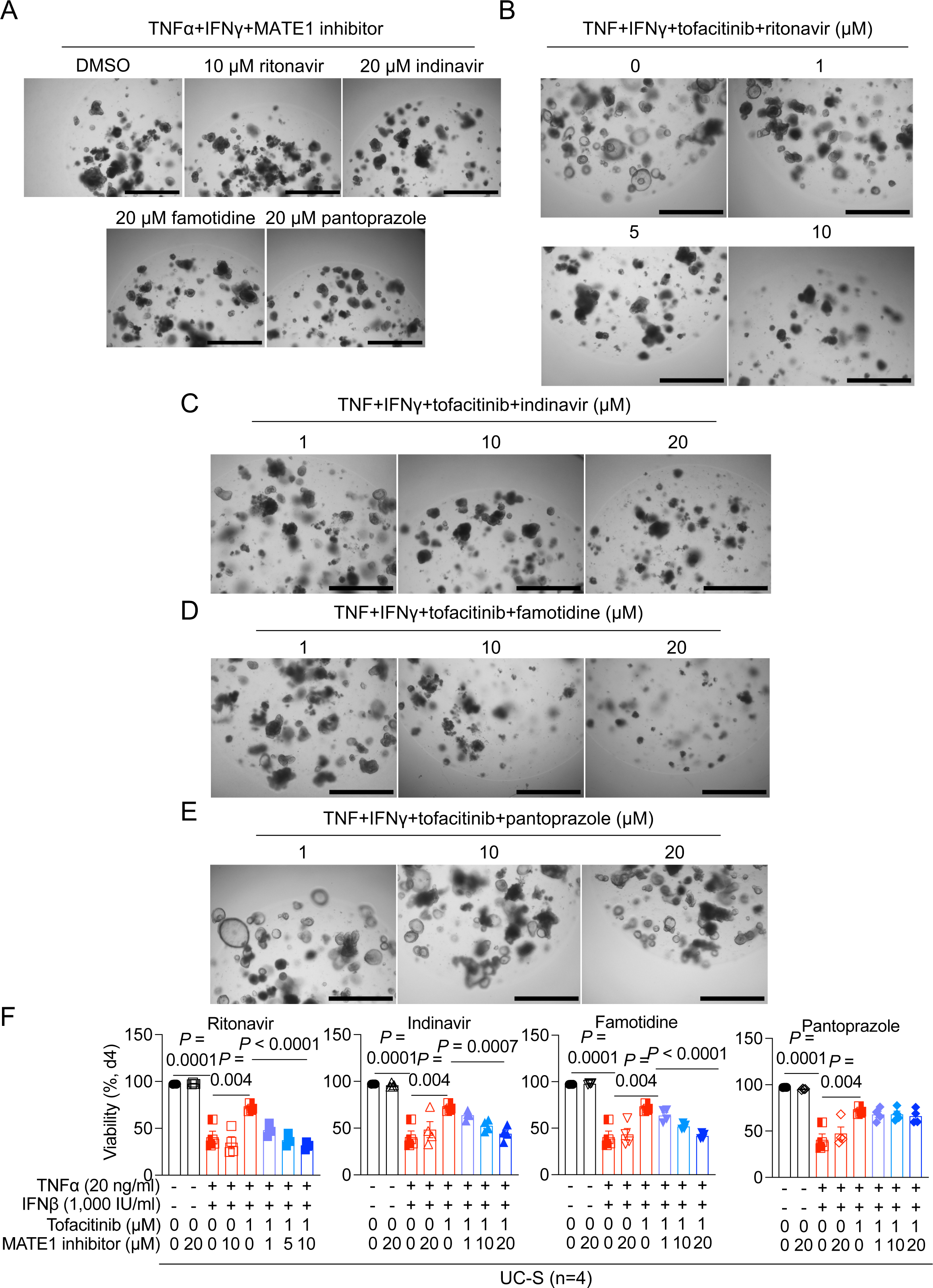
MATE1 inhibitors suppress the ability of tofacitinib to rescue the cell death of UC-S organoids. (A-E) Representative images of UC-S organoids stimulated TNFα+IFNγ±tofacitinib in the presence of 20 μM of MATE1 inhibitors (A), cytokine and tofacitinib co-treated organoids in the presence of increasing concentration of ritonavir (B), indinavir (C), famotidine (D), or pantoprazole (E) on day 4. (F) Quantification of viability of cytokine-stimulated organoids in A-E on day 4. Bars in A-E, 1,000 μm. Concentrations of cytokines, tofacitinib, and MATE1 inhibitors are indicated in F. Concentrations of cytokines in A-E are the same in F. Data points in F represent individual organoid lines and are mean of three technical replicates. Bars represent mean±SEM. Indicated *P* values by unpaired *t* test in F.

**Supplementary Figure 12.**
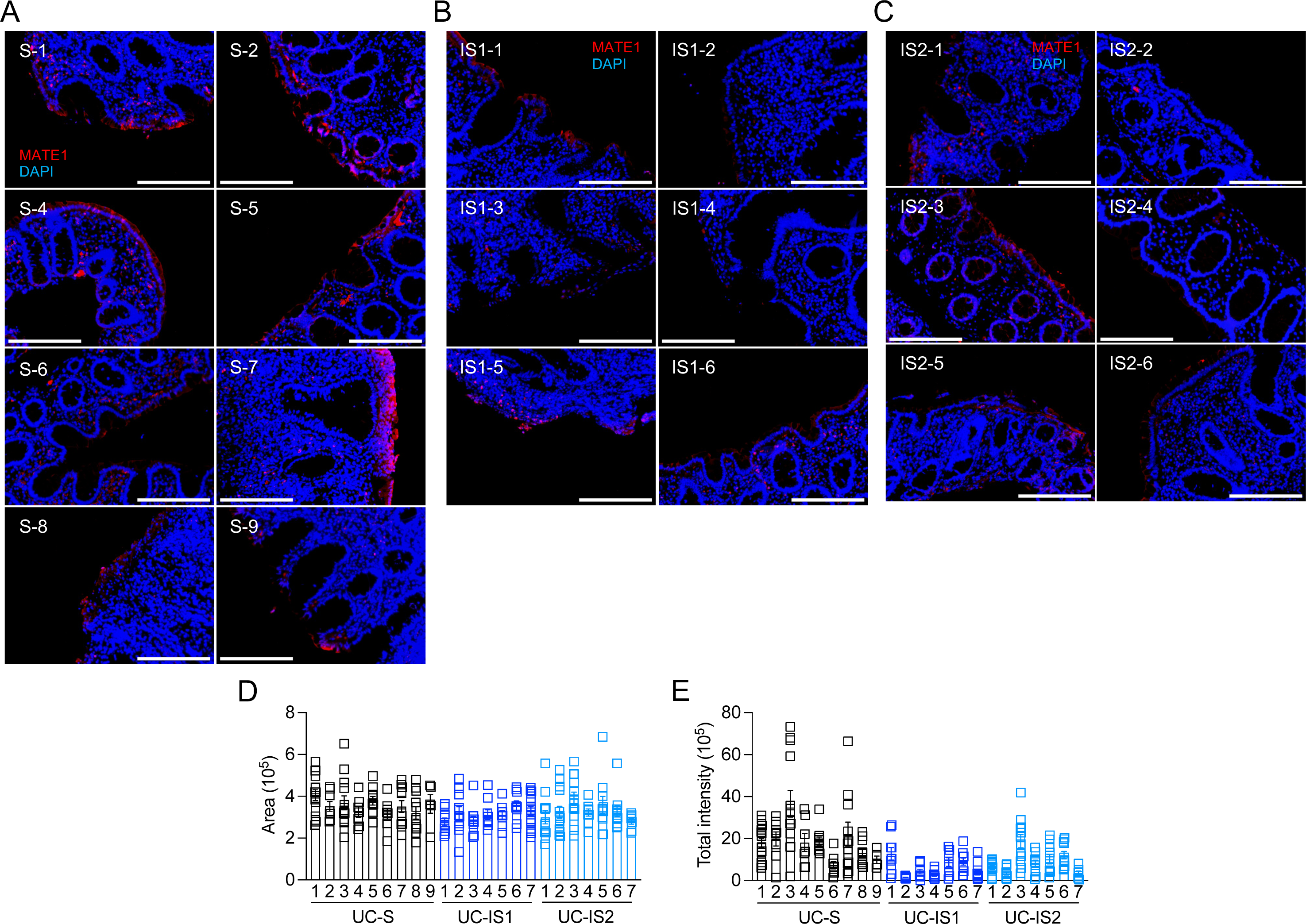
Representative images of MATE1 staining in primary colonic tissue. (A-C) Representative MATE1 staining images in primary tissues of ascending colon from which UC-S (S-1, 2 and 4 to 9, A), UC-IS1 (IS1-1 to 6, B), or UC-IS2 (IS2-1 to 6) organoids were established. Images of remaining organoids are included in Figure 5A. (D and E) Quantification of the total intensity (D) of MATE1 staining and surface area (E) in each visual field from the primary tissues from which the indicated organoid lines were established. Bars in A-C, 200 μM. Data points in D and E are visual field. Bars represent mean±SEM.

**Supplementary Figure 13.**
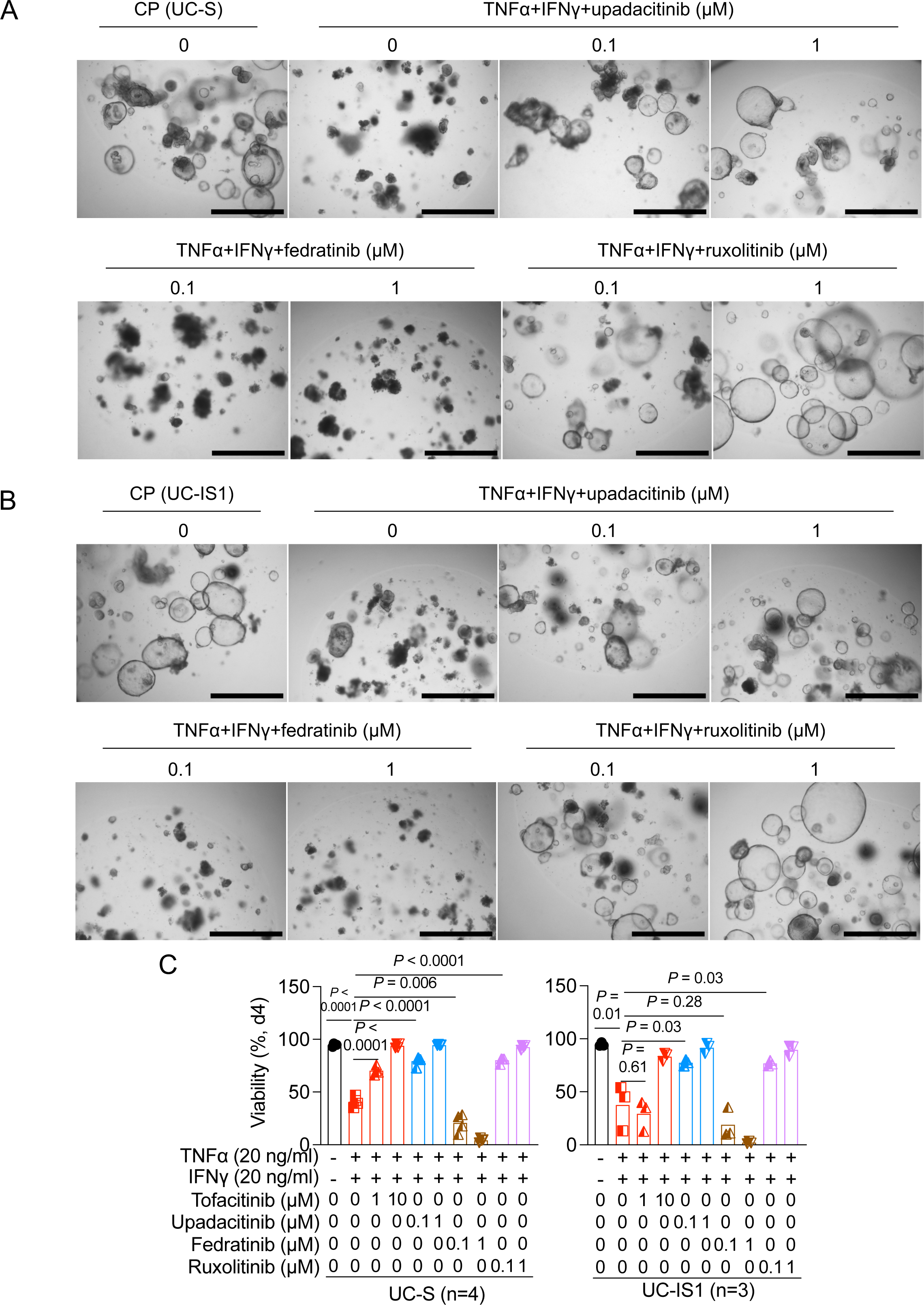
Upadacitinib and ruxolitinib rescue the cell death of cytokine-stimulated UC-IS1. (A and B) Representative images of UC-S (A) or UC-IS1 (B) organoids treated with TNFα+IFNγ in the presence of increasing concentration of upadacitinib, fedratinib, or ruxolitinib on day 4. (C) Quantification of viability of UC-S (left) or UC-IS1 (right) organoids in A and B. Bars in A and B, 1,000 μm. Concentrations of cytokines and JAK inhibitors are indicated in C. Concentrations of cytokines in A and B are the same in C. Data points in C represent individual organoid lines and are mean of three technical replicates. Bars represent mean±SEM. Indicated *P* values by unpaired *t* test in C.

**Supplementary Figure 14.**
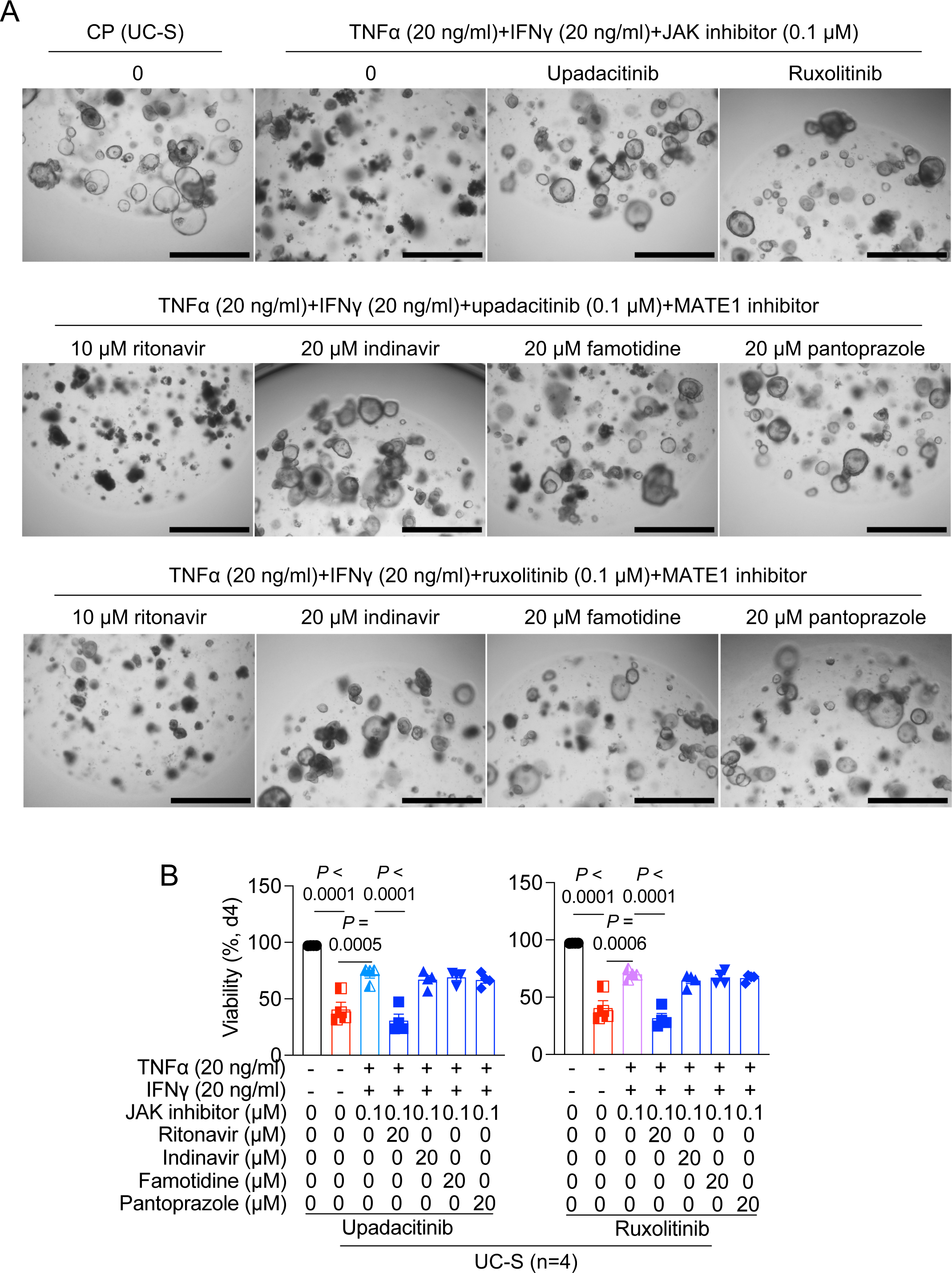
Rescue of cytokine-induced cell death by upadacitinib and ruxolitinib is dependent on OCT1. (A and B) representative images (A) and quantification of viability (B) of upadacitinib-(left) or ruxolitinib-(right) treated UC-S organoids stimulated with TNFα+IFNγ in the presence of ritonavir, indinavir, famotidine, or pantoprazole on day 4. Bars in A, 1,000 μm. Concentrations of cytokines, upadacitinib, and ruxolitinib are indicated in A and B. Data points in A represent individual organoid lines and are mean of two technical replicates. Bars represent mean±SEM. Indicated *P* values by unpaired *t* test in B.

**Supplementary Table 1.** Characteristics of patients on week 0.

| Characteristics on week 0 | UC (n = 48) |  |
| --- | --- | --- |
| Age (yrs) | Mean (range) | 34.7 (17-68) |
| Sex—male | N (%) | 26 (54.2) |
| Disease duration (yrs) | Mean (range) | 9.5 (2-40) |
| **UC disease location** |  |  |
| Proctitis (E1) | N (%) | 1 (2.1) |
| Left sided (E2) | N (%) | 14 (29.2) |
| Pancolitis (E3) | N (%) | 33 (68.5) |
| **Prior anti-TNF therapy use** |  |  |
| 0 | N (%) | 7 (14.6) |
| 1 | N (%) | 29 (60.4) |
| 2 | N (%) | 7 (14.6) |
| ≥3 | N (%) | 2 (4.2) |
| **Prior vedolizumab use** | N (%) | 33 (68.8) |
| **Prior ustekinumab use** | N (%) | 10 (20.8) |
| **Clinical biomarkers** |  |  |
| CRP*, mg/L | Mean (range) | 16.4 (0.1-110) |
| Fecal calprotectin, µg/g | Mean (range) | 3115.5 (78-21169) |
| **Mayo Endoscopic subscore disease activity** |  |  |
| Mayo 1 | N (%) | 1 (2.1) |
| Mayo 2 | N (%) | 13 (27.1) |
| Mayo 3 | N (%) | 34 (70.8) |
| **Concomitant medication** |  |  |
| None | N (%) | 31 (64.6) |
| Systemic corticosteroids | N (%) | 17 (35.4) |

\*CRP, C-reactive protein

**Supplementary Table 2.** Mean total Mayo scores of the patients on week 0 and week 24.

|  | **Week 0** | **Week 24** |
| --- | --- | --- |
| **Total (n = 48)** |  |  |
| Total Mayo | 8.8 | 5.2 |
| Partial Mayo | 6.1 | 3.9 |
| MES* (0-3) | 2.7 | 1.5 |
| **Response (n = 18)** |  |  |
| Total Mayo | 8.6 | 4.8 |
| Partial Mayo | 6.0 | 3.5 |
| MES* (0-3) | 2.6 | 1.3 |
| **Partial / Nonresponse (n = 27)** |  |  |
| Total Mayo | 8.8 | 5.4 |
| Partial Mayo | 6.2 | 4.0 |
| MES* (0-3) | 2.7 | 1.6 |
| **Lost to follow-up (n = 3)** |  |  |
| Total Mayo | 9 | N/A |
| Partial Mayo | 6.4 | N/A |
| MES( 0-3) | 2.6 | N/A |

\*MES, Mayo endoscopic activity

**Supplementary Table 3.** Information for donors. M, male; F, female; UC, ulcerative colitis; HD, healthy donor; UC-S, tofacitinib-sensitive UC patient; UC-IS1, group 1 tofacitinib-insensitive UC patient; UC-IS2, group 2 tofacitinib-insensitive UC patient; W0, week 0; W24, week 24; N/A, not available; RP, responder; PRP, partial responder; NRP, non-responder.

| **Name** | **Sex** | **Disease** | **Organoid**<br>**(group)** | **Total Mayo**<br>**(W0)** | **Total Mayo**<br>**(W24)** | **Tofacitinib**<br>**responsiveness** |
| --- | --- | --- | --- | --- | --- | --- |
| HD1 | F | N/A | O (HD1) | N/A | N/A | N/A |
| HD2 | M | N/A | O (HD2) | N/A | N/A | N/A |
| HD3 | M | N/A | O (HD3) | N/A | N/A | N/A |
| HD4 | M | N/A | O (HD4) | N/A | N/A | N/A |
| P1 | M | UC | O (UC-IS2-2) | 8 | 4 | PRP |
| P2 | M | UC | X | 10 | N/A | NRP |
| P3 | M | UC | O (UC-IS1-5) | 8 | 11 | NRP |
| P4 | F | UC | X | 12 | N/A | N/A |
| P5 | F | UC | O (UC-IS1-6) | 10 | 12 | NRP |
| P6 | F | UC | X | 10 | N/A | NRP |
| P7 | M | UC | X | 10 | 6 | PRP |
| P8 | F | UC | O (UC-S-4) | 5 | 0 | RP |
| P9 | M | UC | O (UC-S-5) | 8 | 2 | RP |
| P10 | F | UC | X | 10 | 1 | RP |
| P11 | F | UC | O (UC-IS2-5) | 8 | 0 | RP |
| P12 | M | UC | O (UC-S-6) | 11 | 6 | NRP |
| P13 | F | UC | O (UC-S-3) | 10 | 1 | RP |
| P14 | M | UC | O (UC-S-8) | 10 | 7 | PRP |
| P15 | F | UC | X | 10 | N/A | NRP |
| P16 | M | UC | X | 9 | 0 | RP |
| P17 | M | UC | X | 9 | 1 | RP |
| P18 | M | UC | X | 7 | 1 | RP |
| P19 | F | UC | X | 6 | 1 | RP |
| P20 | M | UC | X | 6 | 4 | RP |
| P21 | M | UC | X | 11 | N/A | PRP |
| P22 | F | UC | X | 10 | 0 | RP |
| P23 | M | UC | X | 10 | N/A | N/A |
| P24 | F | UC | X | 11 | 1 | PRP |
| P25 | M | UC | X | 8 | 7 | NRP |
| P26 | F | UC | X | 10 | 11 | NRP |
| P27 | F | UC | O (UC-IS2-4) | 7 | 0 | RP |
| P28 | F | UC | O (UC-IS2-5) | 8 | 7 | RP |
| P29 | M | UC | O (UC-IS1-3) | 7 | 11 | NRP |
| P30 | F | UC | X | 8 | 5 | PRP |
| P31 | M | UC | X | 10 | N/A | NRP |
| P32 | F | UC | X | 10 | N/A | N/A |
| P33 | F | UC | O (UC-S-9) | 10 | 0 | RP |
| P34 | F | UC | X | 8 | 8 | PRP |
| P35 | F | UC | O (UC-IS1-4) | 7 | 11 | NRP |
| P36 | M | UC | X | 12 | 11 | NRP |
| P37 | M | UC | X | 7 | 1 | RP |
| P38 | M | UC | O (UC-S-2) | 7 | 1 | RP |
| P39 | M | UC | O (UC-IS2-7) | 10 | 10 | NRP |
| P40 | M | UC | X | 6 | 10 | NRP |
| P41 | F | UC | O (UC-IS1-7) | 11 | 11 | NRP |
| P42 | M | UC | X | 9 | 9 | PRP |
| P43 | F | UC | O (UC-IS1-2) | 8 | 9 | NRP |
| P44 | M | UC | O (UC-IS1-1) | 6 | 5 | PRP |
| P45 | F | UC | O (UC-IS2-1) | 10 | 11 | NRP |
| P46 | M | UC | O (UC-S-1) | 9 | 5 | PRP |
| P47 | F | UC | O (UC-S-7) | 7 | 2 | RP |
| P48 | M | UC | O (UC-IS2-3) | 7 | 0 | RP |

**Supplementary Table 4.**
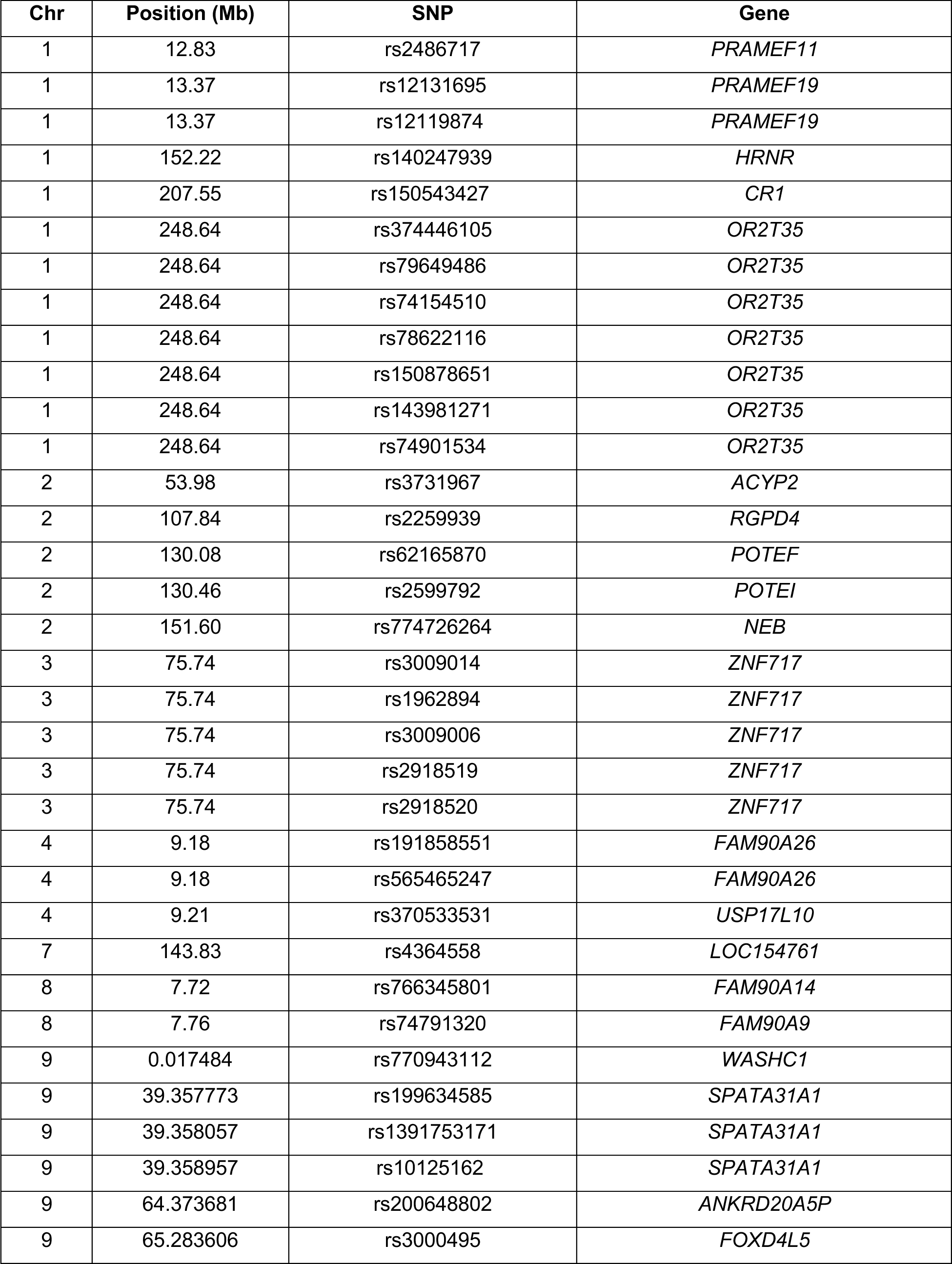
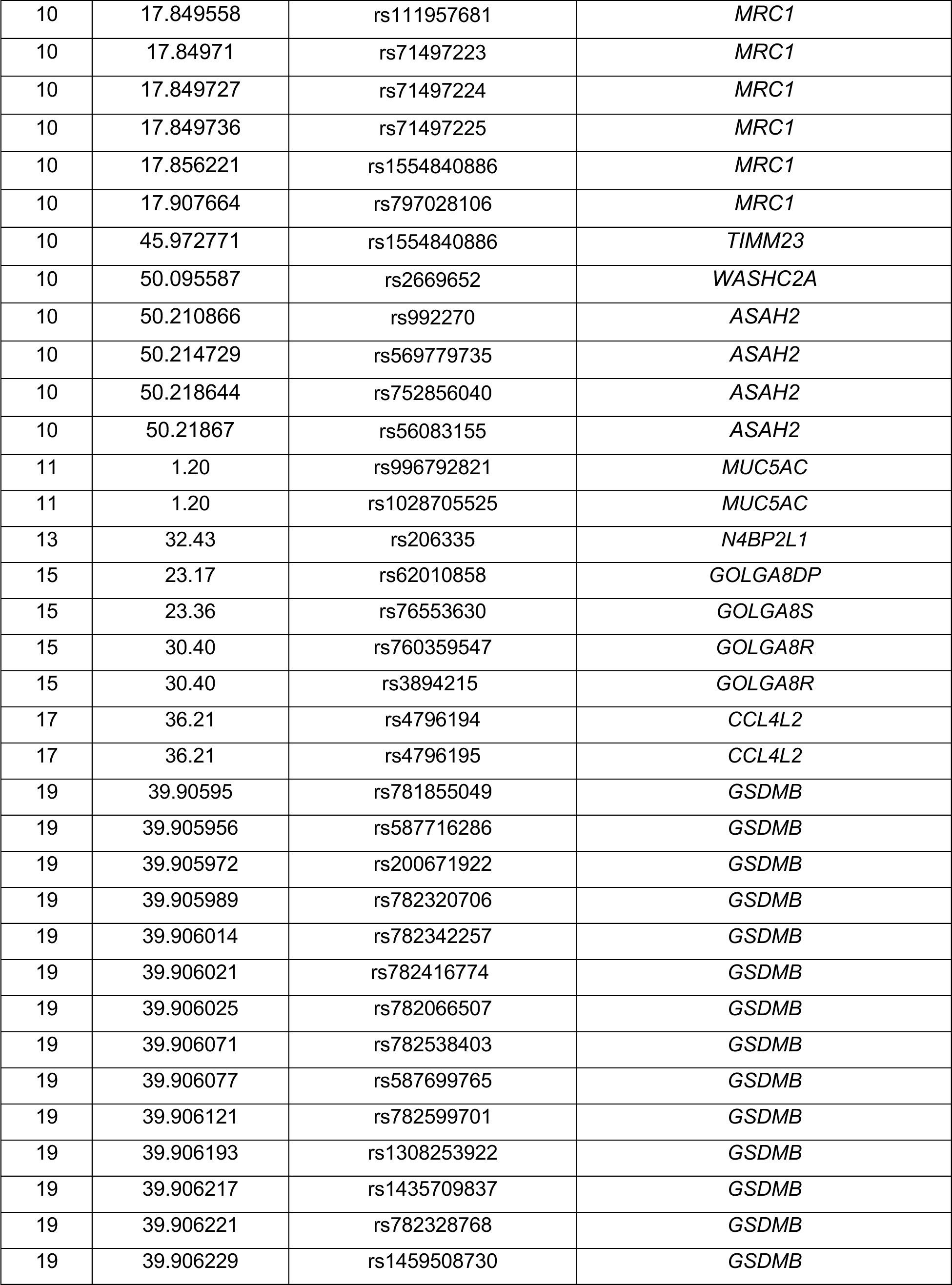
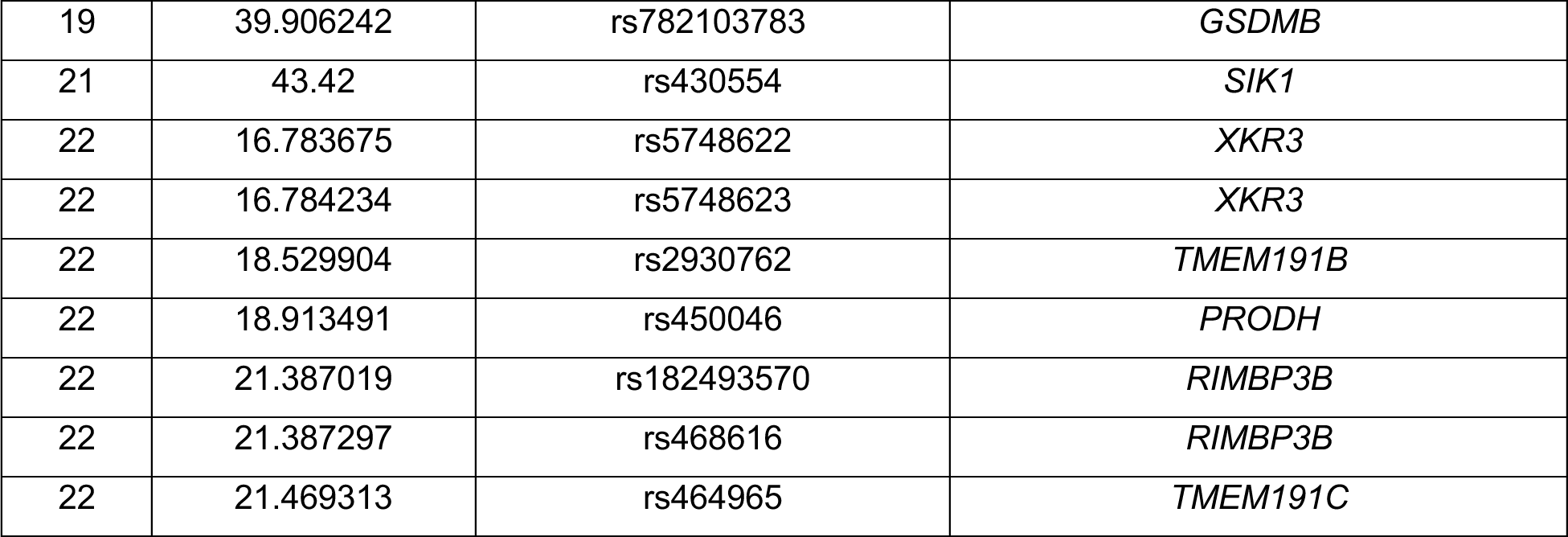
SNPs that are significantly associated with tofacitinib-enrolled patients. The position given is the middle of the locus window, with all positions relative to human reference genome (GRCh38). Chr, chromosome; Mb, megabase; SNP, single nucleotide polymorphism.

